# The impact of purifying and background selection on the inference of population history: problems and prospects

**DOI:** 10.1101/2020.04.28.066365

**Authors:** Parul Johri, Kellen Riall, Hannes Becher, Laurent Excoffier, Brian Charlesworth, Jeffrey D. Jensen

## Abstract

Current procedures for inferring population history generally assume complete neutrality - that is, they neglect both direct selection and the effects of selection on linked sites. We here examine how the presence of direct purifying selection and background selection may bias demographic inference by evaluating two commonly-used methods (MSMC and *fastsimcoal2*), specifically studying how the underlying shape of the distribution of fitness effects (DFE) and the fraction of directly selected sites interact with demographic parameter estimation. The results show that, even after masking functional genomic regions, background selection may cause the mis-inference of population growth under models of both constant population size and decline. This effect is amplified as the strength of purifying selection and the density of directly selected sites increases, as indicated by the distortion of the site frequency spectrum and levels of nucleotide diversity at linked neutral sites. We also show how simulated changes in background selection effects caused by population size changes can be predicted analytically. We propose a potential method for correcting for the mis-inference of population growth caused by selection. By treating the DFE as a nuisance parameter and averaging across all potential realizations, we demonstrate that even directly selected sites can be used to infer demographic histories with reasonable accuracy.

## INTRODUCTION

The characterization of past population size change is a central goal of population genomic analysis, with applications ranging from anthropological to agricultural to clinical (see review by Beichman *et al*. 2018). Furthermore, use of an appropriate demographic model provides a necessary null model for assessing the impact of selection across the genome (*e*.*g*., Teshima *et al*. 2006; Thornton and Jensen 2007; Jensen *et al*. 2019). Multiple strategies have been proposed for performing demographic inference, utilizing expectations related to levels of variation, the site frequency spectrum, linkage disequilibrium, and within- and between-population divergence (*e*.*g*., Gutenkunst et al. 2009; Li and Durbin 2011; Lukic and Hey 2012; Excoffier et al. 2013; Harris and Nielsen 2013; Bhaskar et al. 2015; Boitard et al. 2016; Sheehan and Song 2016; Ragsdale and Gutenkunst 2017; Kelleher et al. 2019; Speidel et al. 2019; Steinrücken et al. 2019).

Although many methods perform well when evaluated under the standard assumption of neutrality, it is difficult in practice to assure that the nucleotide sites used in empirical analyses experience neither direct selection nor the effects of selection at linked sites. For example, inference is often performed using intergenic, 4-fold degenerate, or intronic sites. While there is evidence for weak direct selection on all of these categories in multiple organisms (*e.g*., Haddrill *et al*. 2005; Chamary and Hurst 2005; Andolfatto 2005; Lynch 2007; Zeng and Charlesworth 2010; Choi and Aquadro 2016; Jackson *et al*. 2017), it is also clear that such sites near or in coding regions will also experience background selection (BGS; Charlesworth *et al*. 1993; Charlesworth 2013), and may periodically be affected by selective sweeps as well (Messer and Petrov 2013; Schrider et al. 2016). These effects are known to affect the local underlying effective population size, and alter both the levels and patterns of variation and linkage disequilibrium (Charlesworth et al. 1993; Kaiser and Charlesworth 2009; O’Fallon et al. 2010; Charlesworth 2013; Nicolaisen and Desai 2013; Ewing and Jensen 2016; Johri et al. 2020).

However, commonly-used approaches for performing demographic inference that assume complete neutrality, including *fastsimcoal2* (Excoffier et al. 2013) and MSMC/PSMC (Li and Durbin 2011; Schiffels and Durbin 2014), have yet to be thoroughly evaluated in the light of this assumption, which is likely to be violated in practice. There are, however, some exceptions, as well as subsequent suggestions on how best to choose the least-affected genomic data for analysis (Pouyet et al. 2018). Rather than investigating existing software, Ewing and Jensen (2016) implemented an approximate Bayesian (ABC) approach quantifying the impact of BGS effects, demonstrating that weak purifying selection can generate a skew towards rare alleles that would be mis-interpreted as population growth. Under certain scenarios, this resulted in a many-fold mis-inference of population size change. However, the effects of the density of directly selected sites and the shape of the distribution of fitness effects (DFE), which are probably of great importance, have yet to be fully considered. Spanning the range of these potential parameter values is important for understanding the implications for empirical applications. For example, the proportion of the genome experiencing direct purifying selection can vary greatly between species, with estimates ranging from ∼3-8% in humans, ∼12% in rice, 37-53% in *D. melanogaster*, and 47-68% in *S. cerevisiae* (Siepel et al. 2005; Liang et al. 2018). Furthermore, many organisms have highly compact genomes, with ∼88% of the *E. coli* genome (Blattner et al. 1997), and effectively all of many virus genomes, being functional (*e.g*., >95% of the SARS-CoV-2 genome, Wu *et al*. 2020).

While such estimates allow us to approximate the effects of BGS in some model organisms, in which recombination and mutation rates are well known, it is difficult to predict these effects in the vast majority of study systems. Moreover, while the genome-wide mean of *B*, a widely-used measure of BGS effects that measures the level of variability relative to neutral expectation, can range from ∼0.45 in *D. melanogaster* to ∼0.94 in humans (Charlesworth 2013; but see Pouyet *et al*. 2018), existing demographic inference approaches are usually applied across organisms without considering this important source of differences in levels of bias. Here, we examine the effects of the DFE shape and functional density on two common demographic inference approaches - the multiple sequentially Markovian coalescent (MSMC) and *fastsimcoal2*. Finally, we propose an extension within the approximate Bayesian computation (ABC) framework to address this issue, treating the DFE as a nuisance parameter and demonstrating greatly improved demographic inference even when using directly selected sites alone.

## RESULTS AND DISCUSSION

### Effects of SNP numbers, density and genome size on inference under neutral equilibrium

The accuracy and performance of demographic inference was evaluated using two popular methods, MSMC (Schiffels and Durbin 2014) and *fastsimcoal2* (Excoffier et al. 2013). In order to assess performance, it was first necessary to determine how much genomic information is required to make accurate inference when the assumptions of neutrality are met. Chromosomal segments of varying sizes (1 Mb, 10 Mb, 50 Mb, 200 Mb, and 1 Gb) were simulated under neutrality and demographic equilibrium (*i.e.,* a constant population size of 5000 diploid individuals) with 100 independent replicates each. For each replicate this amounted to the mean [SD] number of segregating sites for each diploid individual being 1,944 [283], 9,996 [418], 40,046 [957] and 200,245 [1887]; for 50 diploid individuals, these values were 10,354 [225], 51,863 [567], 207,118 [1139] and 1,035,393 [2476] for 10 Mb, 50 Mb, 200 Mb and 1 Gb, respectively. Use of MSMC resulted in incorrect inferences for all segments smaller than 1 Gb (Supp Figures 1, 2). Specifically, very strong recent growth was inferred instead of demographic equilibrium, although ancestral population sizes were correctly estimated. In addition, when two or four diploid genomes were used for inference, MSMC again inferred a recent many-fold growth for all segment sizes even when the true model was equilibrium, but performed well when using 1 diploid genome with large segments (Supp Figures 1, 2). These results suggest caution when performing inference with MSMC on smaller regions or genomes, specifically when the number of SNPs is less than ∼200,000 per single diploid individual. Extra caution should be used when interpreting population size changes inferred by MSMC when using more than 1 diploid individual.

When using *fastsimcoal2* to perform demographic inference, parameters were accurately estimated for all chromosomal segment sizes when the correct model (*i.e*., equilibrium) was specified (Supp Table 1). However, when model selection was performed using a choice of four models (equilibrium, instantaneous size change, exponential size change, and instantaneous bottleneck), the correct model was chosen more often (∼30% of replicates) when the simulated chromosome sizes were small (1 and 10 Mb), while an alternative model of either instantaneous size change or instant bottleneck was increasingly preferred for larger regions (Supp Tables 2, 3), although the estimates of ancestral sizes were correct. This finding suggests that the non-independence of SNPs may result in model mis-identification. Indeed, since the model choice procedure assumes that SNPs are independent, the true number of independent SNPs is overestimated, which results in an overestimation in the confidence of the model choice with increasing amount of data. However, it is interesting to note that the parameter values underlying the non-constant size preferred model were often pointing towards a constant-population size (see below). When model selection was performed using sparser SNP densities (*i.e*., 1 SNP per 5 kb, 50 kb or 100 kb), the correct model was recovered for longer chromosomes up to 200 Mb (Supp Tables 2, 3; Supp Figures 3, 4), although model selection was slightly less accurate for smaller chromosomes due to the decrease in the total amount of data. As suspected, the biases introduced by the non-independence of SNPs were found to be concordant with the level of linkage disequilibrium amongst SNPs used for the analysis (for 10 SNP windows, in which SNPs were separated by 50 kb (100 kb), mean *r*^2^ = 0.027 (0.020), compared to the all-SNP mean *r*^2^ of 0.118, and to the completely unlinked SNPs mean *r*^2^ of 0.010; Supp Table 4). Additionally, AIC performed on partially linked SNPs may impose an insufficient penalty on larger number of parameters, resulting in an undesirable preference for parameter-rich models. We found that implementing a more severe penalty improved inference considerably, even for 1 Gb chromosome sizes (Supp Table 5, 6). This model selection performance, the potential corrections related to increased penalties, as well as the total number of SNPs and SNP thinning, should be investigated on a case-by-case basis in empirical applications, owing to the contribution of multiple underlying parameters (*e.g*., chromosome length, recombination rates, and SNP densities).

In the light of this performance assessment, all further analyses were restricted to characterizing demographic inference on data that far exceeded 1 Gb and roughly matched the structure and size of the human genome - for every diploid individual, 22 chromosomes (autosomes) of size 150 Mb each were simulated, which amounted to roughly 3 Gb of total sequence. Ten independent replicates of each parameter combination were performed throughout, and inference utilized one and fifty diploid individuals for MSMC and *fastsimcoal2,* respectively.

### Effect of the strength of purifying selection on demographic inference

In order to test the accuracy of demographic inference in the presence of BGS, all 22 chromosomes were simulated with exons of size 350 bp each, with varying sizes of introns and intergenic regions (see Methods) in order to vary the fraction (5%, 10% and 20%) of the genome under selection. Because the strength of selection acting on deleterious mutations affects the distance over which the effects of BGS extend, demographic inference was evaluated for various DFEs (Table 1). The DFE was modelled as a discrete distribution with four fixed classes: 0 ≤ 2*N_anc_s* < 1, 1 ≤ 2*N_anc_s* < 10, 10 ≤ 2*N_anc_s* < 100 and 100 < 2*N_anc_s* < 2*N_anc_*, where *N_anc_* is the ancestral effective population size and *s* is the reduction in the fitness of the homozygous mutant relative to wildtype. The fitness effects of mutations were uniformly distributed within each bin, and assumed to be semi-dominant, following a multiplicative fitness model for multiple loci; the DFE shape was altered by varying the proportion of mutations belonging to each class, given by *f*_0_, *f*_1_, *f*_2_, and *f*_3_, respectively (see Methods). Three DFEs highly skewed towards a particular class were initially used to assess the impact of the strength of selection on demographic inference (with the remaining mutations equally distributed amongst the other three classes): DFE1: a DFE in which 70% of mutations have weakly deleterious fitness effects (*i.e*., *f*_1_ = 0.7); DFE2: a DFE in which 70% of mutations have moderately deleterious fitness effects (*i.e*., *f*_2_ = 0.7); and DFE3: a DFE in which 70% of mutations have strongly deleterious fitness effects (*i.e*., *f*_3_ = 0.7). A DFE with equal proportions of all deleterious classes (*i.e*., DFE4: *f*_0_= *f*_1_ = *f*_2_ = *f*_3_= 0.25) was also simulated to evaluate the combined effect of different selective strengths. In addition, two bimodal DFEs consisting of only the neutral and the strongly deleterious class of mutations were simulated to characterize the role of strongly deleterious mutations (DFE5: a DFE in which 50% of mutations have strongly deleterious effects (*i.e*., *f*_3_ = 0.5) with the remaining being neutral; and DFE6: a DFE in which 30% of mutations were strongly deleterious (*i.e*., *f*_3_ = 0.3) with the remaining being neutral).

**Table 1:** Proportion (*f*_i_) of mutations in each class of the discrete distribution of fitness effects (DFE) simulated in this study.

| | $f_0$ | $f_1$ | $f_2$ | $f_3$ |
| --- | --- | --- | --- | --- |
| **DFE1** | 0.1 | 0.7 | 0.1 | 0.1 |
| **DFE2** | 0.1 | 0.1 | 0.7 | 0.1 |
| **DFE3** | 0.1 | 0.1 | 0.1 | 0.7 |
| **DFE4** | 0.25 | 0.25 | 0.25 | 0.25 |
| **DFE5** | 0.5 | 0.0 | 0.0 | 0.5 |
| **DFE6** | 0.7 | 0.0 | 0.0 | 0.3 |

In order to understand the effects of BGS, exonic sites were masked, and only linked neutral intergenic and intronic sites were used for demographic inference by both MSMC and *fastsimcoal2* (although comparisons are presented under certain models to analyses based on non-masked datasets). The three demographic models examined were (1) demographic equilibrium, (2) a 30-fold exponential growth, mimicking the recent growth experienced by European human populations, and (3) ∼6-fold instantaneous decline, mimicking the out-of-Africa bottleneck in human populations (Figure 1a). Although these models were parameterized using previous estimates of human demographic history (Supp Table 7; Gutenkunst *et al*. 2009), these basic demographic scenarios are applicable to many organisms, although the magnitudes of population size changes in this case may represent an extreme. Under neutrality, inference of parameters of all three simulated demographic models was highly accurate with both MSMC and *fastsimcoal2* (Figure 1a; Supp Table 8). However, when inferring parameters using *fastsimcoal2*, the time of change in case of the population decline model was consistently over-estimated when SNPs separated by 5 kb were used, while the time was accurately inferred when using all SNPs (Supp Table 8). We therefore present our results using all SNPs throughout (with comparisons to 1 SNP per 5 kb and 1 SNP per 100 kb thinning, under certain models), and recommend caution when implementing thinning procedures.

**Figure 1:**
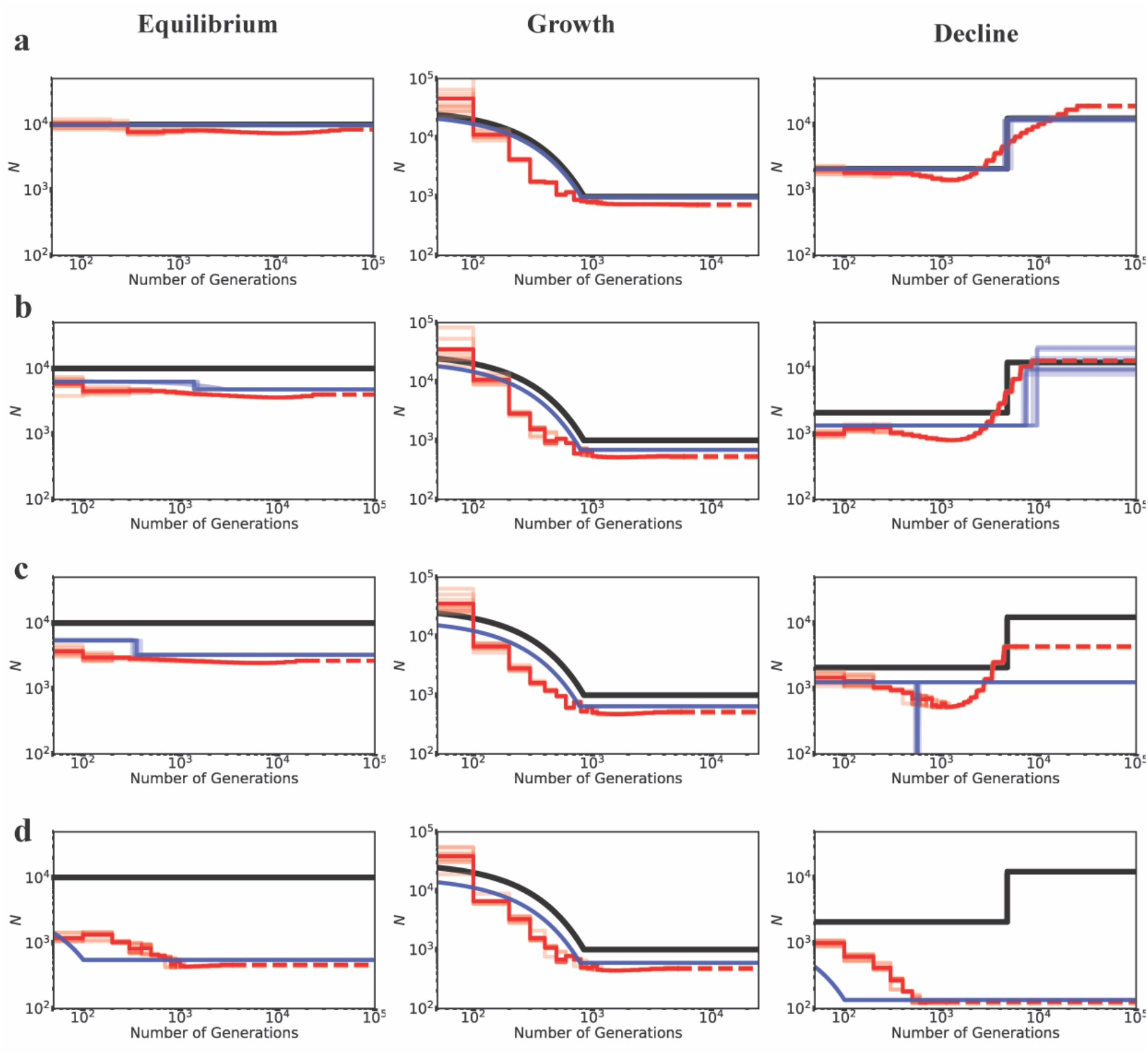
Inference of demography by MSMC (red lines; 10 replicates) and *fastsimcoal2* (blue lines; 10 replicates) with and without BGS, under demographic equilibrium (left column), 30-fold exponential growth (middle column), and ∼6-fold instantaneous decline (right column). The true demographic models are depicted as black lines, with the x-axis origin representing the present day. (a) All genomic sites are strictly neutral. Exonic sites experience purifying selection specified by (b) DFE1, (c) DFE2, and (d) DFE3 (see Table 1). Exons represent 20% of the genome, and exonic sites were masked/excluded when performing demographic inference, quantifying the effects of BGS alone. The dashed lines represent indefinite extensions of the ancestral population sizes. Detailed methods including command lines can be found at: https://github.com/paruljohri/demographic_inference_with_selection/blob/main/CommandLines/Figure1.txt

Under demographic equilibrium, when 20% of the genome experiences direct selection (with masking of the directly selected sites), we found the true population size to be underestimated as expected, and recent population growth mis-inferred (Figure 1, Supp Figure 5), even when only 1 SNP per 100kb was used and a higher AIC penalty was employed (Supp Figure 6). Conversely, when the true demographic model was characterized by a recent 30-fold growth, demographic inference was accurate and performed equally well for both MSMC and *fastsimcoal2*, with the exception of a slight underestimation of the ancestral population size for all DFE types. When the true model was population decline, weakly deleterious mutations alone did not affect inference drastically with either method, and it was possible to recover the true model (*i.e*., decline vs growth) by *fastsimcoal2* in all replicates (Supp Figure 7). However, moderately and strongly deleterious mutations resulted in an underestimation of population size and the inference of an instantaneous bottleneck and strong recent growth respectively, to the extent that population decline was misinterpreted as a bottleneck/growth in all replicates (Supp Figures 5,7). Strong recent growth was inferred (in the presence of moderately and strongly deleterious mutations) even when SNPs separated by 100 kb were used, and an increased penalty was employed against parameter-rich models (Supp Figure 6). We further tested the effect of BGS on demographic inference when changes in population size were less severe, namely, when population growth and decline were only 2-fold, with qualitatively similar results (Supp Figure 8).

Finally, given the strong evidence that most organisms have a bi-modal DFE with a significant proportion of strongly deleterious or lethal mutations (Sanjuán 2010; Jacquier et al. 2013; Kousathanas and Keightley 2013; Bank et al. 2014; Charlesworth 2015; Galtier and Rousselle 2020), we investigated the effect of this strongly deleterious class further. Thus, for comparison with the above, we simulated a rather extreme case in which 30% or 50% of exonic mutations were strongly deleterious with fitness effects uniformly sampled between 100 ≤ 2*N*_anc_*s* ≤ 2*N*_anc_, with the remaining mutations being neutral (*i.e*., DFE5 and DFE6; see Table 1). As with the above results, both equilibrium and decline models were falsely inferred as growth, with an order of magnitude underestimation of the true population size (Figure 2).

**Figure 2:**
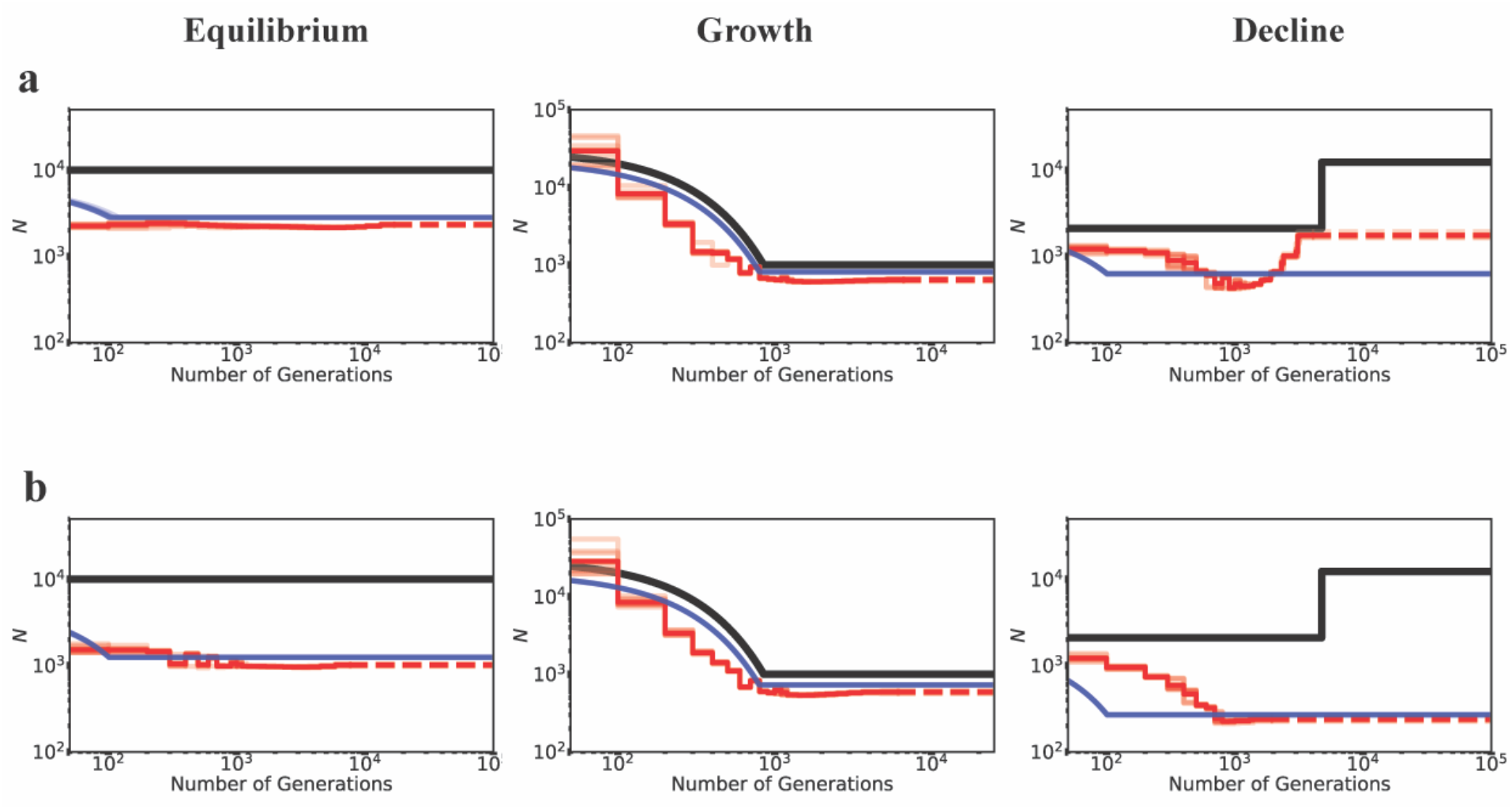
Inference of demography by MSMC (red lines; 10 replicates) and *fastsimcoal2* (blue lines; 10 replicates) in the presence of BGS generated by strongly deleterious mutations. Directly selected sites comprised 20% of the genome and were masked when performing demographic inference. Exons experience purifying selection specified by (a) DFE6 and (b) DFE5 (see Table 1). The true demographic models are given as black lines, with the x-axis origin representing the present day. The dashed lines represent indefinite extensions of the ancestral population sizes.Detailed methods including command lines can be found at: https://github.com/paruljohri/demographic_inference_with_selection/blob/main/CommandLines/Figure2.txt

In sum, neglecting BGS frequently results in the inference of population growth, almost regardless of the true underlying demographic model.

### Effects of density and inclusion/exclusion of directly selected sites on inference

Although we have shown that the presence of purifying selection biases demographic inference, the extent of mis-inference necessarily depends on the fraction of the genome experiencing direct selection. We therefore compared models in which 5%, 10% or 20% of the genome was functional. For this comparison, equal proportions of mutations in each DFE bin were assumed corresponding to DFE4 (Table 1). As before, when the true model was growth, inference was unbiased, with a slight underestimation of ancestral population size when 20% of the genome experienced selection (Figure 3). Population decline was inferred reasonably well if less than 10% of the genome experienced direct selection, but could be mis-inferred as growth with greater functional density, as shown in Figure 3. Similarly, the extent to which population size was under-estimated at demographic equilibrium increased with the fraction of the genome under selection. Finally, it is noteworthy that many changes in population size that were falsely inferred were greater than 2-fold in size, suggesting the need for great caution when inferring such changes from real data.

**Figure 3:**
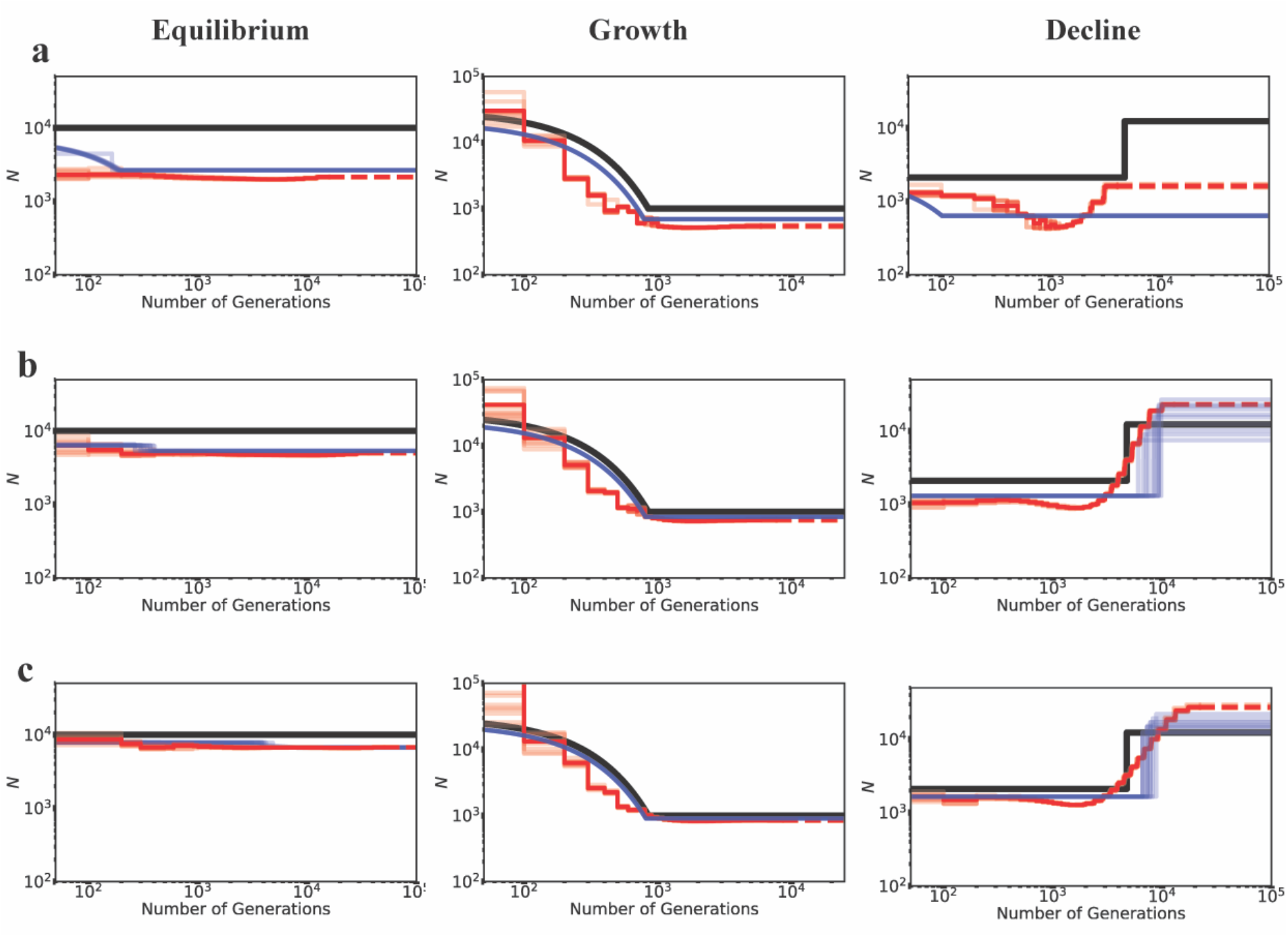
Inference of demography by MSMC (red lines; 10 replicates) and *fastsimcoal2* (blue lines; 10 replicates) in the presence of BGS with varying proportions of the genome under selection, for demographic equilibrium (left column), exponential growth (middle column), and instantaneous decline (right column). Exonic sites were simulated with purifying selection with all *f*_i_ values equal to 0.25 (DFE4; see Table 1), and were masked when performing inference. Directly selected sites comprise (a) 20% of the simulated genome, (b) 10% of the simulated genome, and (c) 5% of the simulated genome. The true demographic models are given by the black lines, with the x-axis origin representing the present day. The dashed lines represent indefinite extensions of the ancestral population sizes. Detailed methods including command lines can be found at: https://github.com/paruljohri/demographic_inference_with_selection/blob/main/CommandLines/Figure3.txt

Importantly, the results presented do not significantly differ between inference performed while including directly selected sites (*i.e*., no masking of functional regions; Supp Figure 9) versus inference performed using linked neutral sites (*i.e*., masking functional regions; Figures 1-3). These results suggest that the exclusion of exonic sites, which is often assumed to provide a sufficiently neutral dataset to enable accurate demographic inference, is not necessarily a satisfactory solution unless gene density is low. For example, demographic inference would naturally be expected to be less biased by BGS for human-like genomes with a relatively low functional density, and more biased in genomes with higher functional density like *D. melanogaster*.

### Effect of BGS on model selection and inferred time of size change using *fastsimcoal2*

In order to quantify the effects of BGS on model selection, four competing models were used for inference: equilibrium, instantaneous size change (growth/decline), exponential size change (growth/decline), and an instantaneous bottleneck. Although demographic equilibrium was almost always inferred as an instantaneous size change (70-100% of replicates), the fitted parameters of the size change model were nearly indistinguishable from the correct model (Figure 1a). In other words, the inferred size change was so inconsequential so as to be nearly a constant-size model, suggesting that parameter estimation is usually more reliable than model selection. When there was a substantial proportion of highly deleterious mutations (DFE3 and DFE5), exponential growth was generally inferred. However, when there was a true size change, *fastsimcoal2* performed well in distinguishing between exponential *vs.* instantaneous change models even in the presence of BGS (Supp Figures 5, 6), provided that the magnitude of size change was large. When size changes were on the order of 2-fold, exponential growth was consistently inferred to be instantaneous.

With respect to model choice between growth and decline in the presence of BGS (irrespective of instantaneous *vs.* exponential change), as the density of selected sites and strength of purifying selection increased, both equilibrium and decline models were more likely to be inferred as growth and occasionally as instantaneous bottlenecks (Supp Figure 7), while true growth models were generally chosen correctly. It should be added that with such large chromosome sizes (3 Gb of total sequence data), model selection was not observed to vary between replicates using *fastsimcoal2* for any given parameter combination. Thus, in the presence of BGS, high-confidence calls of an incorrect underlying demographic model appear likely.

With regard to the time of inferred size change, when the true model was exponential growth, the model was always correctly identified and inference of the time of change was slightly under-estimated in the presence of BGS (Supp Figures 10, 11), consistent with the fact that BGS will further skew the site frequency spectrum towards rare alleles. When the true model was decline, and the model was correctly identified as such, the time of change was modestly over-estimated (Supp Figures 12, 13) – up to ∼2-fold for 6× growth and 2.5-fold for 2× growth (when 20% of the genome was exonic).

### Effect of heterogeneity in recombination rates, mutation rates, and repeat masking

Variation in recombination and mutation rates, as well as the masking of repeat regions, may also affect demographic inference procedures. We evaluated this issue by simulating heterogeneity in both mutation and recombination rates (based on estimated human genome maps, as described in the Methods section), and masking 10% of each simulated segment drawing from the empirical distribution of repeat lengths in the human genome (Supp Figure 14). In general, inferences under neutrality (Supp Figures 15-17) as well as under BGS (Supp Figures 18-20) were not affected to a great extent, suggesting such heterogeneity to have a comparatively minor role for the parameter space considered in this study. Thus, serious mis-inference is more likely to be caused by selection. These observations also suggest that simulations performed with mean rates of recombination and mutation, as in this study, are sufficient to evaluate biases caused by BGS.

### Effects of BGS on diversity and the SFS under various demographic models: theoretical expectations versus simulation results

To better understand how BGS can lead to different biases in the inference of population history, we investigated the extent of BGS effects under all three demographic models, with respect to both the expected diversity in the presence of BGS relative to neutrality (*B*), as well as the shape of the SFS at linked neutral sites. First, we found that *B* differed among demographic scenarios, with much lower values in the case of equilibrium and decline, concordant with stronger demographic mis-inference (Figure 4). After a population decline, *B* was lower than that before the size change; while after population expansion, *B* increased relative to that in the ancestral population, sometimes approaching 1 (Figure 4). This may seem paradoxical, given that the magnitude of the scaled selection coefficient (2*Nes*) decreases with decreasing *Ne* (*i.e.*, the efficacy of purifying selection decreases, and could thus be expected to result in larger values of *B* under population decline). Conversely, with increasing *Ne*, *B* should be expected to reduce.

**Figure 4:**
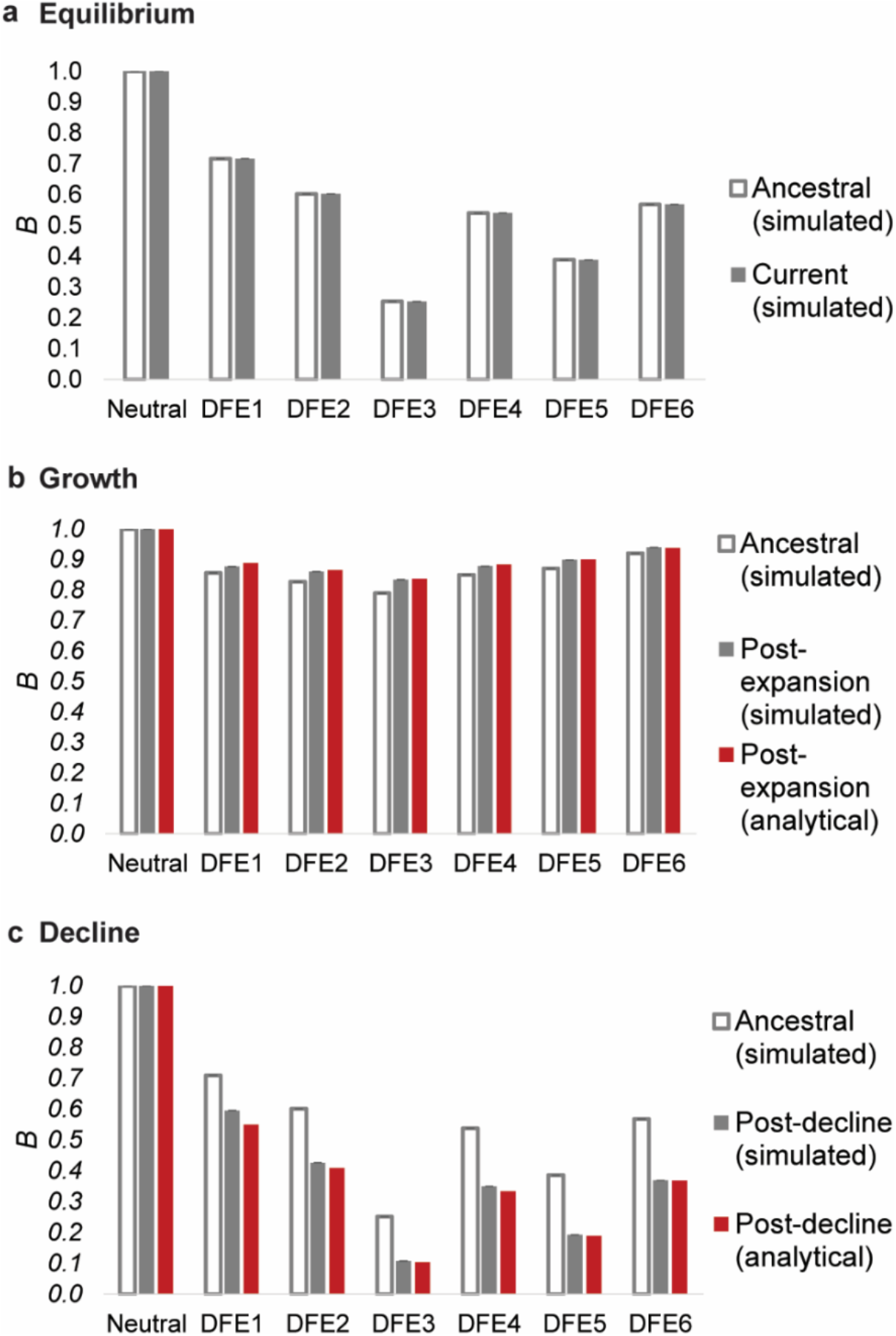
Nucleotide site diversity with BGS (*B*) relative to its purely neutral expectation (*π*_0_) for varying DFEs (specified in Table 1) and demographic scenarios. The results are shown for (a) demographic equilibrium, (b) population growth, and (c) population decline. All cases refer to size changes forward in time, the ancestral *B* (*i.e*., *B* pre-change in population size) is shown in white bars, *B* post-change in population size is shown in solid gray bars, and the analytical expectations for the post-size change *B* is shown as red bars. Exonic sites comprised ∼10% of the genome, roughly mimicking the density of the human genome. Detailed methods including command lines can be found at: https://github.com/paruljohri/demographic_inference_with_selection/blob/main/CommandLines/Figure4.txt

However, these expectations apply only once a population has maintained a given *Ne* for sufficient time such that mutation-drift-equilibrium has been approached. During the initial stages of population size change, and shortly afterwards, the dynamics of *B* tend to show a trend opposing these long-term expectations (see also Figure 5 of Torres *et al*. 2020). This is because differences in *Ne* caused by different initial levels of BGS cause differences in the rates of response to changes in population size – a small value of *Ne* (corresponding to low *B*) results in a faster response compared with a high value (Fay and Wu 1999; Hey and Harris 1999; Pool and Nielsen 2007; Pool and Nielsen 2009; Campos et al. 2014; Torres et al. 2020). In other words, diversity in a growing population will increase more rapidly in regions experiencing stronger BGS than in completely neutral regions, while diversity in a declining population will decrease at a faster rate in regions with BGS relative to those with neutrality, resulting in temporarily higher and lower *B*, respectively. The relative diversity values observed with different initial equilibrium *B* values after a short period of population size change may thus be very different from both the initial and final equilibrium values. The overall effect is that there is an apparent increase in *B* immediately following a population decline, and a decrease immediately following an expansion. Analytical models describing these effects are presented in the Appendix. These models used the simulated values of *B* at equilibrium before the population size changes to predict the apparent *B* values at the ends of the periods of size change (see the Methods and Appendix). It can be seen from Figure 4 that there is good agreement between these predictions and the simulation results.

**Figure 5:**
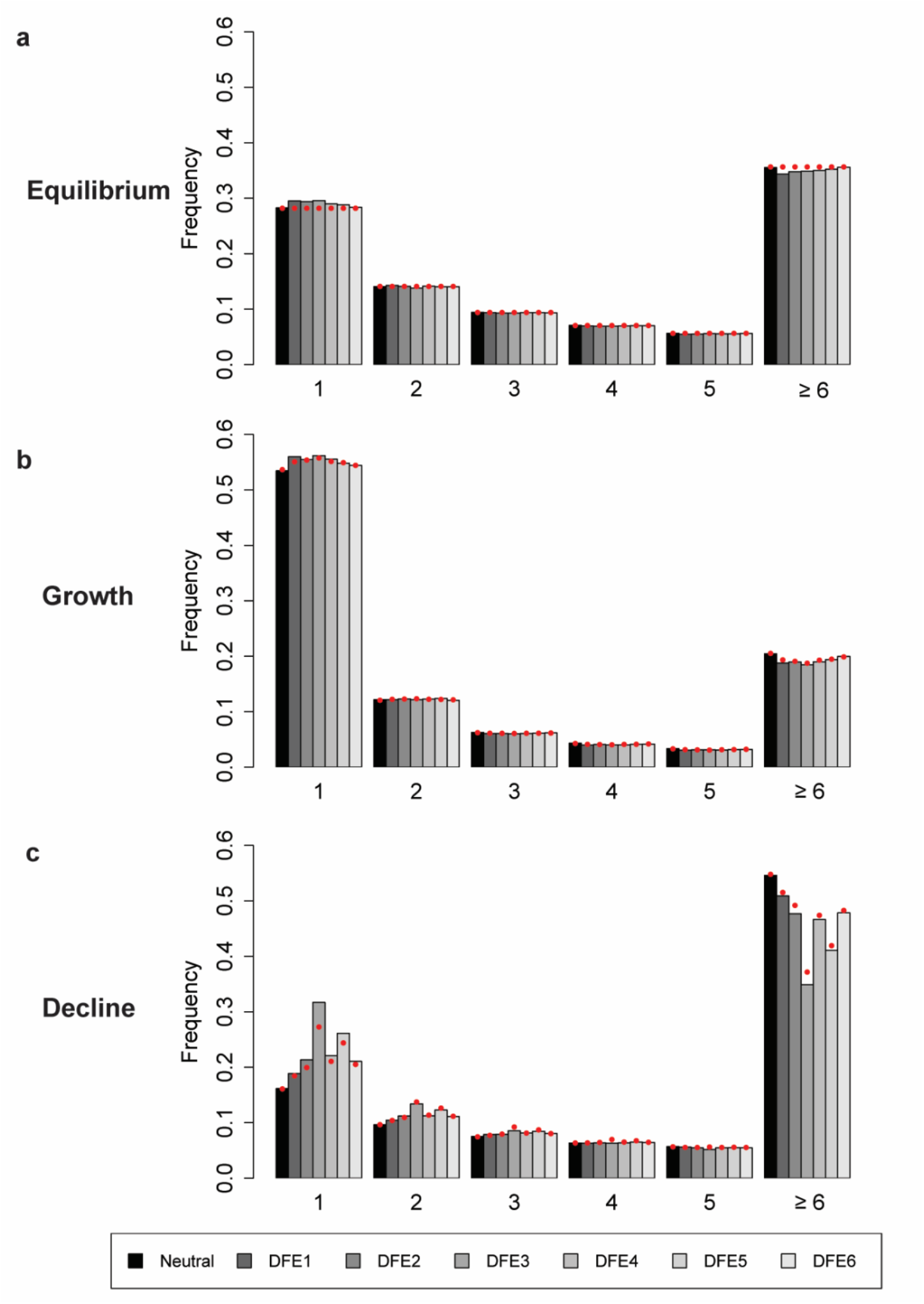
The site frequency spectrum (SFS) of derived allele frequencies at neutral sites from 10 diploid genomes under (a) demographic equilibrium, (b) population growth, and (c) population decline, under the same DFEs as shown in Figure 4. The x-axis indicates the number of sample alleles (out of 20) carrying the derived variant. Exonic sites comprised ∼10% of the genome, roughly mimicking the density of the human genome. The red solid circles give the values predicted analytically with a purely neutral model, but correcting for BGS by using the *B* values of the ancestral population (*i.e*., pre-change in population size) obtained from simulations, in order to quantify the effective population size. Detailed methods including command lines can be found at: https://github.com/paruljohri/demographic_inference_with_selection/blob/main/CommandLines/Figure5.txt

Because several demographic estimation methods are based on fitting a demographic model to the SFS, it is also of interest to determine whether BGS can skew the SFS to different extents under different demographic models. Although it is well known that BGS causes a skew of the SFS towards rare variants under equilibrium models (Charlesworth et al. 1995; Nicolaisen and Desai 2013), the effect of BGS on the SFS with population size change has not been much explored (but see Johri et al. 2020; Torres et al. 2020). As shown in Figure 5, with a population size decline, the SFS of derived alleles is more skewed towards rare variants when BGS is operating, especially when *B* is initially small, since the effects of BGS work in opposition to the effects of the population size reduction. This difference in the left skew of the SFS with and without BGS is much less noticeable in the case of population expansion, since here the effects of BGS and the expansion act in a similar direction.

As with the estimates of the apparent *B* values discussed above, analytical predictions of the expected SFS after an instantaneous / exponential change in population size can be made, using the values of *B* and the SFS at equilibrium in the ancestral population before the population size change using the formulae of Polanski and Kimmel (2003) and Polanski *et al*. (2003) for the purely neutral case, as described in the Methods section. Importantly, the use of the *B* parameter does not in itself cause a skew in the SFS, it merely affects overall diversity values. Figure 5 shows that the overall shape of the SFS is predicted reasonably well by the analytical results, although deviations are to be expected for the rare allele classes, which are the most sensitive to demographic change and selection. Overall, the results imply that BGS is more likely to bias demographic inference post-decline compared with post-expansion, consistent with the performance of the methods described above. Although it is notable that the SFS can be reasonably well predicted by correcting for the re-scaling effects of BGS if the effects of BGS in the ancestral population are accurately known, the exact allele frequency patterns observed will depend on the timing of population size changes relative to the time of sampling, as well as the value of *B* prior to the size change. The patterns described here thus represent only a small subset of the possibilities.

### A potential solution: averaging across all possible DFEs

As shown above, demographic inference can be strongly affected by BGS effects that have not been taken into account, as well as by direct purifying selection. A potential solution is thus to correct for these effects when performing inferences of population history. A widely-used approach to estimating direct selection effects, DFE-alpha, takes a stepwise approach to inferring demography, by using a presumed neutral class (synonymous sites); conditional on that demography, it then estimates the parameters of the DFE (Keightley and Eyre-Walker 2007; Eyre-Walker and Keightley 2009; Schneider et al. 2011; Kousathanas and Keightley 2013). However, this approach does not include the possibility of effects of selection at linked sites, which can result an over-estimate of population growth, and while the DFE may not be mis-inferred strongly (Kim et al. 2017), there is substantial mis-inference of the DFE if synonymous sites experience direct selection (Johri et al. 2020).

Building on this idea, Johri et al. (2020) recently proposed an approach that includes both direct and background effects of purifying selection, and simultaneously infers the deleterious DFE and demography. By utilizing the decay of BGS effects around functional regions, they demonstrated high accuracy under the simple demographic models examined. Moreover, the method makes no assumptions about the neutrality of synonymous sites, and can thus be used to estimate selection acting on these sites, as well as in non-coding functional elements. However, this computationally-intensive approach is specifically concerned with jointly inferring the DFE and demographic parameters. As such, if an unbiased characterization of the population history is the sole aim, this procedure may be needlessly involved. We thus here examine the possibility of instead treating the DFE as an unknown nuisance parameter, averaging across all possible DFE shapes, in order to assess whether demographic inference may be improved simply by correcting for these selection effects without inferring their underlying parameter values. This approach utilizes functional (*i.e*., directly selected) regions, a potential advantage in populations for which only coding data may be available (*e.g*., exome-capture data; see Jones and Good 2016), or more generally in organisms with largely functional genomes.

In order to illustrate this approach, a functional genomic element was simulated under demographic equilibrium, 2-fold exponential population growth and 2-fold exponential population decline with four different DFE shapes (as described previously, and shown in Figure 6). A number of summary statistics were calculated (see Methods) for the entire functional region. Inference was first performed assuming strict neutrality, and inferring a one-epoch size change (thus estimating the ancestral (*N*_anc_) and current population sizes (*N*_cur_)). As was found with the other inference approaches examined, population sizes were underestimated and a false inference of population growth was observed in almost all cases when selective effects are ignored (Figure 6).

**Figure 6:**
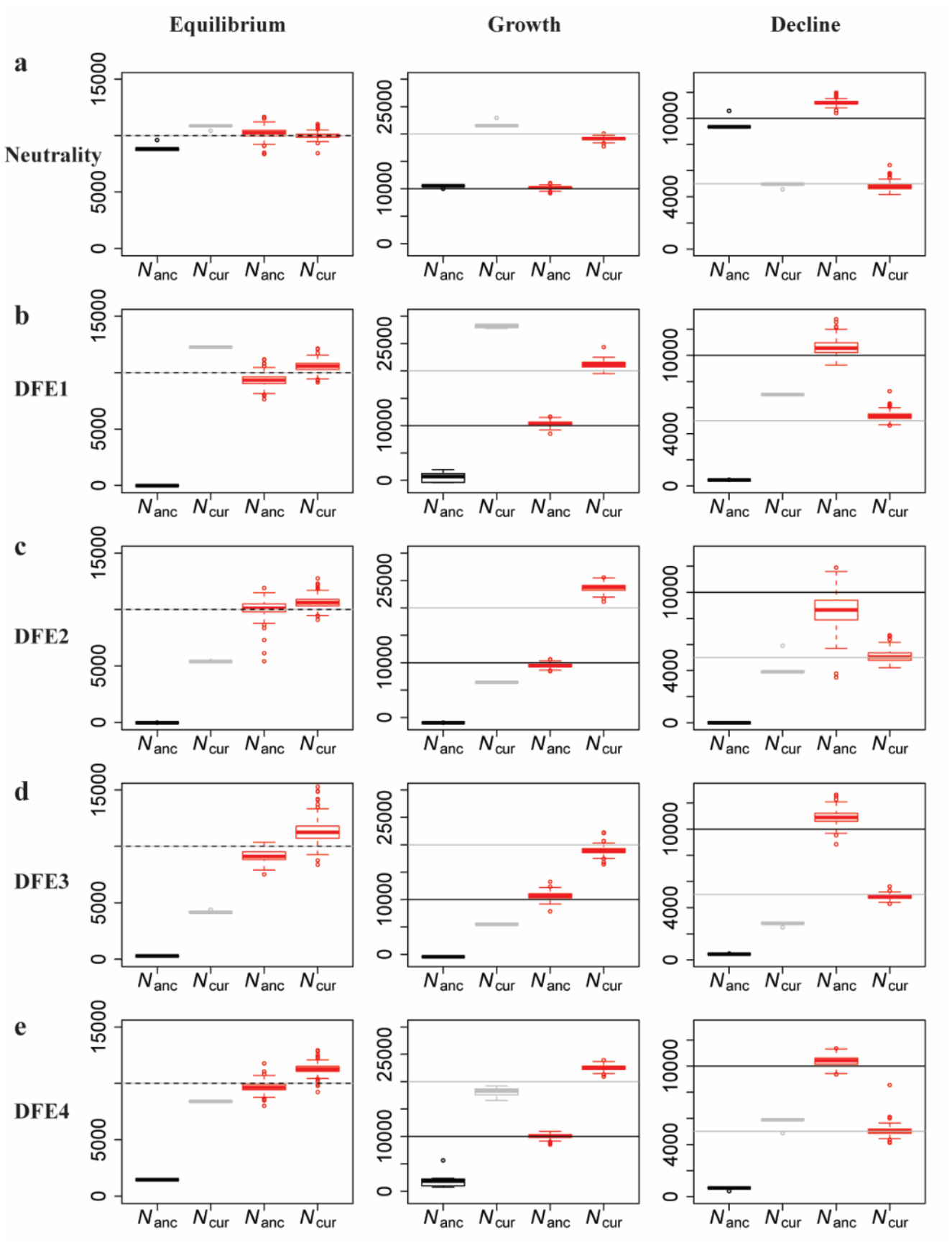
Comparison of estimates of ancestral (*N*_anc_) and current (*N*_cur_) population sizes when assuming neutrality *vs* when varying the DFE shape as a nuisance parameter, using an ABC framework. Inference is shown for demographic equilibrium (left column), 2-fold exponential growth (middle column), and 2-fold population decline (right column), for five separate DFE shapes that define the extent of direct purifying selection acting on the genomic segment for which demographic inference is performed: (a) neutrality, (b) DFE1, (c) DFE2, (d) DFE3, and (e) DFE4 (see Table 1). In each case, the horizontal lines give the true values (black for *N*_anc_; and gray for *N*_cur_) and the box-plots give the estimated values. Black and gray boxes represent estimates when assuming neutrality, while red boxes represent estimates when the DFE is treated as a nuisance parameter. Detailed methods including command lines can be found at: https://github.com/paruljohri/demographic_inference_with_selection/blob/main/CommandLines/Figure6.txt

Next, the assumption of neutrality was relaxed, and mutations were simulated with fitness effects characterized by a discrete DFE, with the fitness classes used above (*f*_0_, *f*_1_, *f*_2_, *f*_3_). Values for *f*_i_ were drawn from a uniform prior between 0 and 1, such that ∑*f*_i_ = 1. Note that no assumptions were made about which sites in the genomic region were functionally important, or regarding the presence/absence of a neutral class. These directly selected sites were then used to infer demographic parameters. We found that, by varying the shape of the DFE, averaging across all realizations, and only estimating parameters related to population history, highly accurate inference of modern and ancestral population sizes is possible (Figure 6). These results demonstrate that, even if the true DFE of a population is unknown (as will always be the case in reality), it is possible to infer demographic history with reasonable accuracy by approximately correcting for these selective effects.

This proposed method is most applicable to organisms in which recombination rates are reasonably well known. If the assumed recombination rate is 2-fold lower than the true rate, the ABC approach infers growth by over-estimating the current population size; correspondingly, if the assumed recombination rate is higher than the true rate, the current population size is under-estimated (Supp Figure 21). Interestingly, in both cases the ancestral population sizes are correctly inferred, consistent with previous results (Johri *et al*. 2020).

## CONCLUSIONS

While commonly used approaches for inferring demography assume neutrality and independence among segregating sites, these assumptions are likely to be violated in practice. In addition, there is considerable evidence for wide-spread effects of selection at linked sites in many commonly studied organisms (Hernandez et al. 2011; Cutter and Payseur 2013; Williamson et al. 2014; Elyashiv et al. 2016; Campos et al. 2017; Booker and Keightley 2018; Pouyet et al. 2018; Ragsdale et al. 2018; Torres et al. 2018; Castellano et al. 2020). Accordingly, we have explored how violations of the assumption of neutrality may affect demographic inference, particularly with regard to the underlying strength of purifying selection and the genomic density of directly selected sites. Generally speaking, the neglect of these effects (*i.e.*, background selection) results in an inference of population growth, with the severity of the growth model roughly scaling with selection strength and density, as well as the inference of historical bottlenecks with some frequency. Thus, when the true underlying model is in fact growth, demographic mis-inference is not particularly severe; when the true underlying model is constant size or decline, the mis-inference can be extreme, with a many-fold underestimation of population size.

However, given that BGS will lead to the false inference of recent growth nearly regardless of the true history, it would be difficult in practice to determine the accuracy of this model without independent information on any given empirical application. Moreover, as the two very different methods investigated here result in highly similar mis-inference, we propose that this performance is unlikely to be a feature of these specific approaches, but rather a quantification of the fact that the underlying genealogies are distorted in the presence of BGS. Thus, these problems are likely to be common to all demographic inference based on polymorphism data.

It is important to note that BGS effects extend over genomic distances in a way that is positively related to the strength of purifying selection. For instance, strongly and moderately deleterious mutations affect patterns of diversity at large genomic distances, whereas mildly deleterious mutations primarily skew allele frequencies at adjacent sites. Thus, if intergenic regions further away from exons are used to perform demographic inference, it is predominantly moderately deleterious mutations that are likely to bias inferences; if these are relatively rare, they may not cause significant problems. In contrast, if synonymous sites are used to infer demographic history, mildly deleterious mutations arising in the coding sequences to which they belong may have significant effects. As we have focused here on relatively sparsely-coding genomes (with human-like gene densities) and used intergenic sites for inference, moderately deleterious mutations resulted in more severe mis-inference. The effect of the decay with distance of BGS due to mildly deleterious mutations depends on multiple parameters. For instance, with an exon of length 500 bp and *Drosophila*-like parameters (*e.g., Ne* = 10^6^; recombination rate = mutation rate = 10^-8^ / site / generation), *B* increases from 0.53 (at 10 bases from the end of the exon) to 0.94 at a distance of 1000 bases. On the other hand, with human-like parameters (*Ne* = 10^4^; recombination rate = mutation rate = 10^-8^ / site / generation) the corresponding change in *B* is only from 0.981 to 0.982 (Supp Table 9).

Thus, mildly deleterious mutations have drastically different effects, depending on the underlying population parameters. While these results certainly suggest that demographic inference ought to be less biased by BGS in neutral regions very distant from functional elements (for species with sufficiently high recombination / functionally sparse genomes), it is noteworthy that purifying selection on moderately and strongly deleterious mutations can have long-range effects, and that the complex interaction of population history with purifying / background selection necessitates a consideration of this topic in any given empirical application.

Comparing the two inference methods investigated here, it appears that *fastsimcoal2* is less prone to inferring false fluctuations in population size. However, both methods falsely infer growth in the presence of BGS, with increasing severity as the density of coding regions increases. The times of population growth inferred by both methods appear to be affected in unpredictable ways when the inferred model is incorrect. When the general model is correctly identified, BGS leads to inference of more recent growth, and more ancient decline, than the reality. In addition, although variation in mutation and recombination rates across the genome alone did not strongly affect demographic inference, our evaluations in the current study are restricted to a specific parameter space resembling those of human populations. The effects of this variation on organisms with more extreme rate fluctuations remain in need of investigation.

It is noteworthy that, even when all sites are strictly neutral or only 5% of the genome experiences direct selection, demographic equilibrium is mis-estimated by MSMC as a series of size changes. The pattern of these erroneous size changes lend a characteristic shape to the MSMC curve (*i.e*., ancient decline and recent growth) which appears to resemble the demographic history previously inferred for the Yoruban population (Schiffels and Durbin 2014), including the time at which changes in population size occurred (Supp Figure 22). Previous work has demonstrated that the resulting demographic model does not in fact fit the observed SFS in the Yoruban population (Beichman et al. 2017; Lapierre et al. 2017). A similar shape has also been inferred in the vervet subspecies (Warren *et al*. 2015; Figure 4), in passenger pigeons (Hung *et al*. 2014; Figure 2), in elephants (Palkopoulou *et al*. 2018; Figure 4), in *Arabidopsis* (Fulgione *et al*. 2018; Figure 3), and in grapevines (Zhou *et al*. 2017; Figure 2A).

Although the inferred population size fluctuations under simulated neutrality are only ∼1.2-fold, in most empirical applications the fluctuations are of a somewhat larger magnitude (∼ 2-fold in pigeons, *Arabidopsis*, and grapevines). Nonetheless, this performance of MSMC under neutral demographic equilibrium is concerning, and adds to the other previously published cautions concerning the interpretation of MSMC results. For example, Mazet *et al*. (2016) and Chikhi *et al*. (2018) demonstrated that, under constant population size with hidden structure, MSMC may suggest false size changes (see also Orozco-terWengel 2016). In addition, MSMC has been reported to falsely infer growth prior to instantaneous bottlenecks (Bunnefeld et al. 2015). In addition, we observed that, if insufficient genomic data are used, or more than one diploid genome is used to perform inference, MSMC falsely infers recent growth of varying magnitudes, the latter having been previously observed by Beichman et al. (2017) and Adrion et al. (2020).

In sum, we find that the effects of purifying and background selection result in similar demographic mis-inference across approaches, and that masking functional sites does not yield accurate parameter estimates. In order to side-step many of these difficulties, our proposed approach of inferring demography by averaging selection effects across all possible DFE shapes within an ABC framework appears to be promising. Utilizing only functional regions, we found a great improvement in accuracy, without making any assumptions regarding the true underlying shape of the DFE or the neutrality of particular classes of sites. As such, this approach represents a more computationally efficient avenue if only demographic parameters are of interest, and ought to be particularly useful in the great majority of organisms in which independent neutral sites either do not exist, or are difficult to identify and verify.

## METHODS

### Simulations of chromosomal segments under neutral equilibrium

When assessing the amount of genomic information required for accurate demographic inference, chromosomal segments of varying sizes (1 Mb, 10 Mb, 50 Mb, 200 Mb and 1 Gb) were simulated under neutral equilibrium. In all cases, the effective population size (*N_e_*) simulated was 5000, and mutation and recombination rates were both 1 × 10^-8^ per site per generation. Simulations were performed with both SLiM 3.1 (Haller and Messer 2019) for a 10*N_e_* generation burn-in, and with msprime 0.7.3 (Kelleher et al. 2016). In all cases 100 replicates were simulated, with the exception of 1 Gb chromosomes simulated by SLiM, in which only 10 replicates were obtained.

### Simulations of human-like chromosomes (with and without selection)

Simulations were performed using SLiM 3.1 (Haller and Messer 2019) for a burn-in of 10*N_anc_* generations, with 10 replicates per evolutionary scenario. For every replicate, 22 chromosomes of 150Mb each were simulated, totaling ∼3 Gb of information per individual genome (similar to the amount of information in a human genome). Within each chromosome, 3 different types of regions were simulated, representing non-coding intergenic, intronic, and exonic regions. Based on the NCBI RefSeq human genome annotation, downloaded from the UCSC genome browser for hg19 (http://genome.ucsc.edu/; Kent *et al*. 2002), mean values of exon sizes and intron numbers per gene were calculated. To represent mean values for the human genome (Lander et al. 2001), each gene comprised 8 exons and 7 introns, and exon lengths were fixed at 350 bp. By varying the lengths of the intergenic and intronic regions, three different genomic configurations with varying densities of functional elements were simulated and compared - with 5%, 10% and 20% of the genome being under direct selection - hereafter referred to as genome5, genome10, and genome20, respectively. Genome5 was comprised of introns of 3000 bp and intergenic sequence of 31000 bp, genome10 of introns of 1500 bp and intergenic sequence of 15750 bp, while genome20 was comprised of introns of 600 bp and intergenic sequence of 6300 bp. The total chromosome sizes of these genomes were approximately 150 Mb (150,018,599 bp, 150,029,949 bp, and 150,003,699 bp) with 2737, 5164, and 11278 genes per chromosome in genome5, genome10, and genome20, respectively. In order to be conservative with respect to the performance of existing demographic estimators, intronic and intergenic regions were assumed to be neutral.

Recombination and mutation rates were assumed to be equal to 1 x 10^-8^ /site / generation. Neither crossover interference nor gene conversion were modeled (see the discussion in Campos and Charlesworth 2019). Exonic regions in the genomes experienced direct purifying selection given by a discrete DFE comprised of 4 fixed classes (Johri et al. 2020), whose frequencies are denoted by *f*_i_: *f0*, with 0 ≤ 2*N*_e_*s* < 1 (*i.e*., effectively neutral mutations), *f1*, with 1 ≤ 2*N*_e_*s* < 10 (*i.e*., weakly deleterious mutations), *f*_2_, with 10 ≤ 2*N*_e_*s* < 100 (*i.e*., moderately deleterious mutations), and *f*_3_, with 100 ≤ 2*N*_e_*s* < 2*N*_e_ (*i.e*., strongly deleterious mutations), where *N*_e_ is the effective population size and *s* is the reduction in fitness of the mutant homozygote relative to wild-type. Within each bin, the distribution of *s* was assumed to be uniform. All mutations were assumed to be semi-dominant. In all cases, the *N*_e_ corresponding to the DFE refers to the ancestral effective population size.

Six different types of DFE were simulated, described by the parameters provided in Table 1. Three different demographic models were tested for each of these DFEs (Supp Table 7): 1) demographic equilibrium, 2) recent exponential 30-fold growth, resembling that estimated for the human CEU population (Gutenkunst et al. 2009), and 3) ∼6-fold instantaneous decline, resembling the out-of-Africa bottleneck in humans (Gutenkunst et al. 2009). For simulations of demographic equilibrium and decline, population sizes and time of change were scaled down by a factor of 10 (with corresponding scaling of the recombination rate, mutation rate, and selection coefficients), while simulations of growth were not scaled.

### Running MSMC

In order to quantify the effect of purifying selection on demographic inference, we used entire chromosomes generated by SLiM to generate input files for MSMC. For comparison, and in order to quantify the effect of BGS alone on demographic inference, we masked the exonic regions to generate input files. For all parameters, MSMC was performed on a single diploid genome, as the results for this case were the most accurate (Supp Figure 1, 2). Input files were made using the script ms2multihetsep.py provided in the msmc-tools-Repository downloaded from https://github.com/stschiff/msmc-tools. MSMC1 and 2 were run as follows: *msmc_1.1.0_linux64bit -t 5 -r 1.0 -o output_genomeID input_chr1.tab input_chr2.tab … input_chr22.tab.* Population sizes obtained from MSMC were plotted up to the maximum number of generations obtained from MSMC, and the final value of the ancestral population size was extended indefinitely as a dashed line.

### Running fastsimcoal2

Inference was performed by masking all exonic SNPs and using all intronic and intergenic SNPs in order to obtain the most accurate estimates. In order to minimize the effects of linkage disequilibrium (LD), SNPs separated by 5 kb or 100 kb were also used for inference in some cases to assess the impact of violating the assumption of independence. When choosing SNPs separated by a particular distance, the first SNP from each chromosome was chosen and if the distance to the next consecutive SNP was greater than or equal to 5 kb/100 kb, that SNP was included, otherwise the next downstream SNP was evaluated. Site frequency spectra (SFS) were obtained for all sets of SNPs for all 10 replicates of every combination of demographic history and DFE. SNPs from all 22 chromosomes were pooled together to calculate the SFS. In the case of SNPs separated by 5 kb/100 kb, the “0” class of the SFS was scaled down by the same extent as the decrease in the total number of SNPs. *Fastsimcoal2* was used to fit each SFS to 4 distinct models: (a) equilibrium, which estimates only a single population size parameter (*N*); (b) instantaneous size change (decline/growth), which fits 3 parameters - ancestral population size (*N_anc_*), current population size (*N_cur_*), and time of change (*T*); (c) exponential size change (decline/growth), which also estimates 3 parameters - *N_anc_*, *N_cur_* and *T*; and (d) an instantaneous bottleneck model with 3 parameters – *N_anc_*, intensity, and time of bottleneck. The parameter search ranges for both ancestral and current population sizes in all cases were specified to be uniformly distributed between 100-500000 individuals, while the parameter range for time of change was specified to be uniform between 100-10000 generations in all models. The intensity of the bottleneck was sampled from a log-uniform distribution between 10^-5^ and 2. The following command line was used to run *fastsimcoal2*:

*fsc26 -t demographic_model.tpl -n 150000 -d -e demographic_model.est -M -L 50 -q,*

Model selection was performed as recommended by Excoffier *et al*. (2013). For each demographic model, the maximum of maximum likelihoods from all replicates was used to calculate the Akaike Information Criterion (AIC) = 2 × number of parameters – 2 × ln(likelihood) = 2 × number of parameters – 2 × ln(10) × *L*_10_, where *L*_10_ is the logarithm (with respect to base 10) of the best likelihood provided by *fastsimcoal2*. For model choice comparison, we also implemented a stricter penalty of 25× (see Supp Table 5, 6), in which case AIC = 25 × number of parameters – 2 × ln(likelihood). The relative likelihoods (Akaike’s weight of evidence) in favor of the *i*^th^ model were then calculated as:

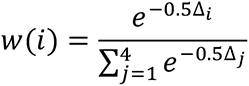

where Δ*_i_* = *AIC_i_* − *AIC*_min_. The model with the highest relative likelihood was selected as the best model, and the parameters estimated using that model were used to plot the final inferred demography.

### Simulations of variable recombination and mutation rates, and repeat masking

In order to simulate variation in recombination and mutation rates, all 22 chromosomes were simulated by mimicking chromosome 6 (∼171Mb) of the human genome. Recombination rates (HapMap) obtained from Yoruban populations (McVean et al. 2004; Myers et al. 2005) were obtained from the UCSC genome browser, while the mutation rate map (https://molgenis26.target.rug.nl/downloads/gonl_public/mutation_rate_map/release2/) was assumed to correspond to estimates obtained from *de novo* mutations (Francioli et al. 2015), as in Castellano *et al*. (2020). Absolute values of mutation rates were normalized in order to maintain the mean mutation rate across the genome at ∼ 1.0 × 10^-8^ per site per generation. Recombination and mutation rate estimates were taken from positions of approximately 10 Mb to 160 Mb, with the recombination map starting at 10010063 bp and the mutation map starting at 10010001 bp. Regions with missing data for either of the two estimates were simulated with rates corresponding to the previous window, except for the case of centromeres in which no recombination was assumed. In order to understand the effect of excluding centromeric regions in empirical studies, the 4Mb region corresponding to the centromere was masked, corresponding to 48.5 to 52.5 Mb of the simulated 150Mb chromosomes. In order to evaluate the effect of masking repeat regions, random segments comprising 10% of each chromosome were masked. The lengths of these segments were drawn from the lengths of repeat regions found in the human genome (Supp Figure 14), as obtained from the repeat regions in the *hg19* assembly of the human genome from the UCSC genome browser.

### Performing inference by approximate Bayesian computation (ABC)

ABC was performed using the R package “abc” (Csilléry et al. 2010), and non-linear regression aided by a neural net (used with default parameters as provided by the package) was used to correct for the relationship between parameters and statistics (Johri et al. 2020). To infer posterior estimates, a tolerance of 0.1 was applied (*i.e*., 10% of the total number of simulations were accepted by ABC in order to estimate the posterior probability of each parameter). The weighted medians of the posterior estimates for each parameter were used as point estimates. ABC inference was performed under two conditions: (1) complete neutrality, or (2) the presence of direct purifying selection. In both cases only 2 parameters were inferred - ancestral (*N*_anc_) and current (*N*_cur_) population sizes. However, in scenario 2, the shape of the DFE was also varied. Specifically, the parameters *f*_0_, *f*_1_, *f*_2_, and *f*_3_ were treated as nuisance parameters and were sampled such that 0 ≤ *f*_i_ ≤ 1, and Σi *f*_i_ = 1, for *i* = 0 to 3. In addition, in order to limit the computational complexity involved in the ABC framework, values of *f*_i_ were restricted to multiples of 0.05 (*i.e*., *f*_i_ ɛ {0.0, 0.05, 0.10, …, 0.95, 1.0} ∀ *i*), which allowed us to sample 1,771 different DFE realizations. Simulations were performed with functional genomic regions, and the demographic model was characterized by 1-epoch changes in which the population either grows or declines exponentially from ancestral to current size, beginning at a fixed time in the past.

For the purpose of illustration, and for a contrast with the human-like parameter set above, parameters for ABC testing were selected to resemble those of *D. melanogaster* African populations. Priors on ancestral and current population sizes were drawn from a uniform distribution between 10^5^-10^7^ diploid individuals, while the time of change was fixed at 10^6^ (∼*N_e_*) generations. In order to simulate functional regions, 94 single-exon genes, as described in Johri *et al*. (2020) and provided in https://github.com/paruljohri/BGS_Demography_DFE/blob/master/DPGP3_data.zip, were simulated with recombination rates specific to those exons (https://petrov.stanford.edu/cgi-bin/recombination-rates_updateR5.pl) (Fiston-Lavier et al. 2010; Comeron et al. 2012). Mutation rates were assumed to be fixed at 3 × 10^-9^ per site per generation (Keightley et al. 2009; Keightley et al. 2014).

All parameters were scaled by the factor 320 in order to decrease computational time, using the principle first described by Hill and Robertson (1966), and subsequently employed by others (Comeron and Kreitman 2002; Hoggart et al. 2007; Kaiser and Charlesworth 2009; Kim and Wiehe 2009; Uricchio and Hernandez 2014; Campos and Charlesworth 2019). The scaled population sizes thus ranged between ∼300–30000 and were reported as scaled values in the main text. One thousand replicate simulations were performed for every parameter combination (*N*_anc_, *N*_cur_, *f*_0_, *f*_1_, *f*_2_, *f*_3_); for performing ABC inference, 50 diploid genomes were randomly sampled without replacement, and summary statistics were calculated using pylibseq 0.2.3 (Thornton 2003). The following summary statistics were calculated across the entire exonic region for every exon: nucleotide site diversity (*π*), Watterson’s *θ*, Tajima’s *D*, Fay and Wu’s *H* (both absolute and normalized), number of singletons, haplotype diversity, LD-based statistics (*r*^2^, *D*, *D*’), and divergence (*i.e*., number of fixed mutations per site per generation after the burn-in period). Means and variances (between exons) of all of the above (a total of 22) were used as final summary statistics to perform ABC. As opposed to the above examples, in this inference scheme only exonic data (*i.e*., directly selected sites) were utilized. Test datasets were generated in exactly the same fashion as described above.

### Analytical expectations for the relative site frequencies

To compute the expected relative frequencies of site frequency classes, the approach of Polanski and Kimmel (2003) was followed. They describe a method for computing the “probability that a SNP has *b* mutant bases”, which is equivalent to the expected site frequency spectrum (SFS) of derived variants. This method (their equations 3-10) allows for the specification of arbitrary population size histories and sample sizes. For reasons of computational precision, a sample size of 10 diploid genomes was chosen. The demographic scenarios were implemented as piecewise functions of the effective population size (counting haploid genomes), and the effect of BGS was included by scaling these functions by values of *B* before population size change as obtained from the forwards-in-time simulations described above. A Mathematica notebook detailing these results is available online (see data availability statement). In addition, analytical expressions can be obtained for pairwise diversity values when there are step changes or exponential growth in population size, as described in the Appendix and in an example program that calculates diversity values after exponential growth.

### Data availability

The following data are publicly available: (1) The general workflow for simulating and performing demographic inference; (2) Scripts used for performing simulations, masking genomes, calculating the SFS, performing model selection and plotting the final results; (3) Input files used to run *fastsimcoal2,* including for the calculated SFS; (4) Output files of MSMC and *fastsimcoal2* for all simulated scenarios; (5) A Mathematica (version 12.1) notebook detailing the calculations of analytical expectations for the relative SFS; (6) An example program (Fortran script) demonstrating how to obtain analytical expressions for values of *B* after exponential growth. All supplemental files, scripts, command lines, and descriptions may be found at: https://github.com/paruljohri/demographic_inference_with_selection

## ACKNOWLEDGEMENTS

We would like to thank Susanne Pfeifer for helpful discussions related to this project, and for feedback on the manuscript. This research was conducted using resources provided by Research Computing at Arizona State University (http://www.researchcomputing.asu.edu) and the Open Science Grid, which is supported by the National Science Foundation and the U.S. Department of Energy’s Office of Science. This work was funded by National Institutes of Health grants R01GM135899 and 1R35GM139383-01 to JDJ.

## APPENDIX

There are two scenarios of population size change for which simple explicit expressions for the expected pairwise coalescent time or diversity can be obtained, without using the methodology of Polanski and Kimmel (2003) and Polanski *et al*. (2003) – a step change in *N* or an exponential growth in *N*. First consider the coalescent process for a step change, where the current and initial effective population sizes are denoted by *N*_e1_ and *N*_e0_, respectively. Let *B* be the background selection parameter at the start of the process of change, corresponding to effective size *N*_e0_. For convenience, time is scaled in units of 2*N*_e1_ generations, and the time of the change in population size on this scale is denoted by*T*_0_, counting back from the present time, *T* = 0. *T*_0_ is assumed to be sufficiently small that *B* remains approximately constant during the period since the change in size. Denote the ratio *N*_e0_/*N*_e1_ by *R.* The derivation for the case of a step change in population size is similar to that given by Pool and Nielsen (2009) for the purpose of comparing X chromosomes and autosomes.

Between times *T* and*T*_0_, coalescence occurs at a rate *B*^−1^ on the chosen timescale, so that the contribution from this period to the net coalescent time for a pair of alleles sampled at *T* = 0 is:

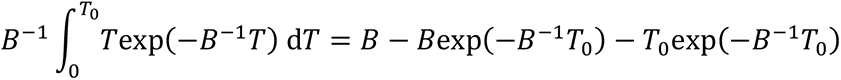

There is a probability of exp(–*B*^−1^*T*_0_) that there is no coalescence when *T* lies between 0 and *T*_0_, after which coalescence occurs at a rate 1/*BR*, giving a net contribution to the coalescence time of:

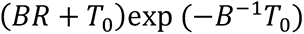

The net coalescence time for the stepwise change with BGS is given by the sum of these two expressions:

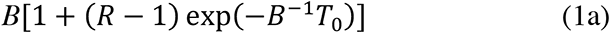

If this expression is compared to the corresponding equation with *B* = 1, the apparent value of *B* at the time of sampling of the pair of alleles is given by:

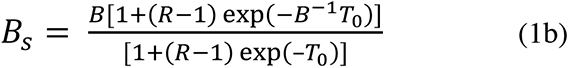

Next, consider a process of exponential change in population size, starting at an initial effective size of *N*_e0_ at *t*_0_ generations in the past and ending at size *N*_e1_, such that the instantaneous growth rate *r* per generation is *r* = ln(*N*_e1_/*N*_e0_)/*t*_0_. The effective population size at time *t* in the past is *N*_e_(*t*) = *N*_e1_exp(–*rt*); with BGS, the rate of coalescence at time *t* is 1/*BN*_e_(*t*). As before, the BGS parameter is assumed to remain constant over the period of population size change. It follows that the probability of no coalescence by generation *t* in the past is:

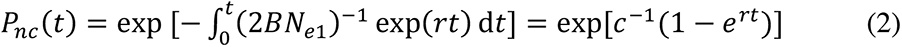

where *c* = 2*BNe*_1_*r*.

The pre-growth period with *t* > *t*_0_ contributes an expected coalescent time of (2*BN_e_*_0_+ *t*_0_)*P_nc_*(*t*_0_), on the scale of generations.

Following Slatkin and Hudson (1991), to obtain the contribution from the period with *t* > *t*_0_, it is convenient to measure time as *τ* = *rt.* The probability of coalescence between *τ* and *τ* + d*τ* is then given by:

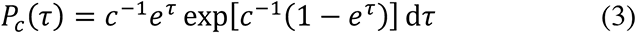

The contribution from this period to the expected coalescent time is given by the integral of *τ Pc*(*τ*) between 0 and *τ*_0_. Following Slatkin and Hudson (1991), by transforming to *u* = exp(*τ*), this contribution can be expressed as the following integral:

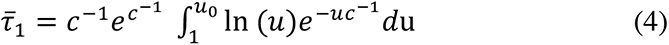

This integral can easily be evaluated numerically. The corresponding mean coalescent time on the scale of generations is obtained by division by *r*, and the result can be added to (2*BN_e_*_0_+ *t*_0_)*P*_nc_(*t*_0_), yielding the net expected coalescent time. By dividing the resulting expression by the corresponding expression with *B* = 1, the apparent BGS effect at the time of sampling can be obtained, in the same way as for the step change model.

## Supplementary Information

**Supp Table 1:**
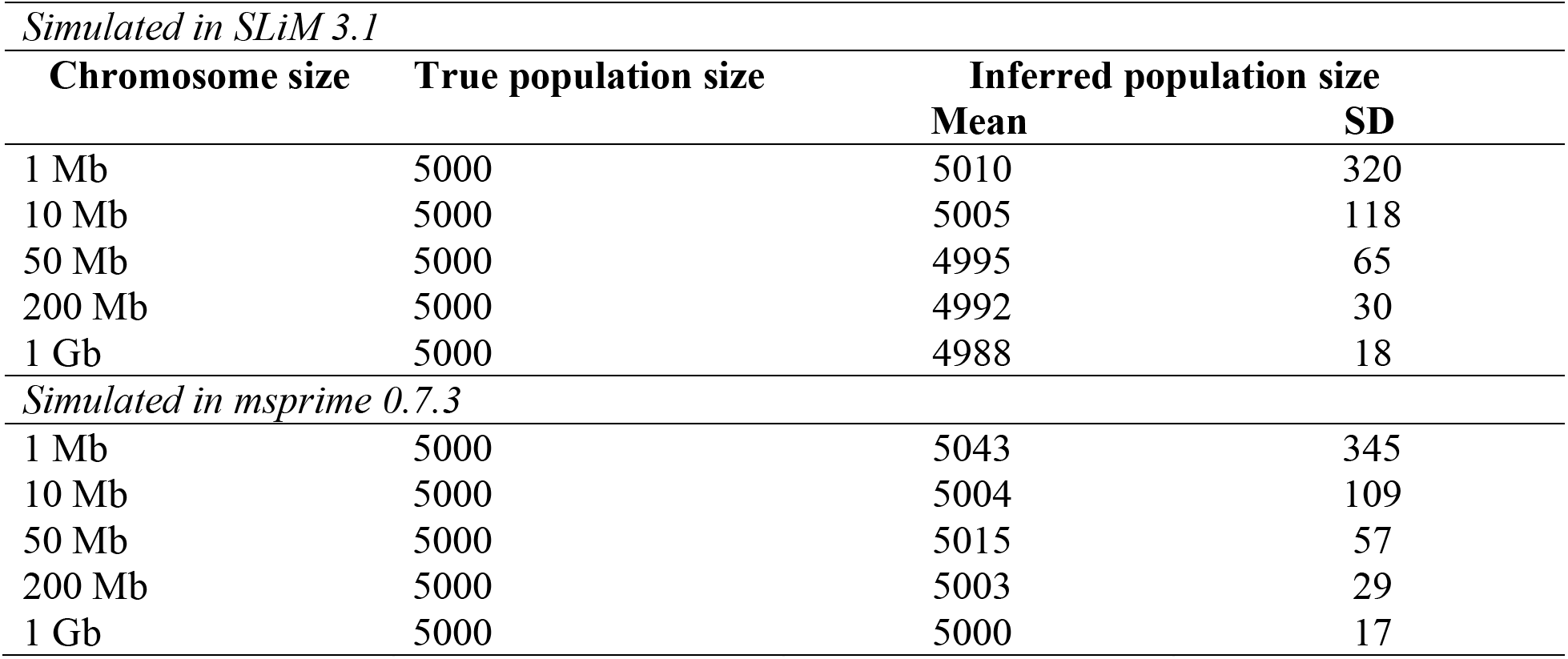
Performance of *fastsimcoal2* under neutrality and demographic equilibrium with varying chromosome sizes, when the correct model was specified. Inference was performed using 50 diploid individuals and all SNPs, and a comparison is given between simulations performed in SLiM (3.1) as well as msprime (0.7.3). The number of replicates for each chromosome size was set to 100, except for 1Gb chromosomes simulated in SLiM for which we report 10 replicates. Detailed methods including command lines can be found here: https://github.com/paruljohri/demographic_inference_with_selection/blob/main/CommandLines/SuppTable1.txt.

**Supp Table 2:** Model selection in *fastsimcoal2* for 100 replicates, in which the true model was neutral equilibrium and selection was performed with 4 models: equilibrium, instantaneous size change, exponential size change, and instantaneous bottleneck. Varying densities of SNPs (all SNPs, 1 per 5 kb, 1 per 50 kb and 1 per 100 kb) were used to perform inference. Simulations were performed using SLiM (3.1) for 10*N* generations. Detailed methods including command lines can be found here: https://github.com/paruljohri/demographic_inference_with_selection/blob/main/CommandLines/SuppFigure3_SuppTable2_5.txt.

| Density of SNPs | Genome length | Equilibrium | Exponential change | Instantaneous change | Instantaneous Bottleneck | Total number of replicates |
| --- | --- | --- | --- | --- | --- | --- |
| all | 1Mb | 33 | 11 | 18 | 38 | 100 |
|  | 10Mb | 32 | 17 | 27 | 24 | 100 |
|  | 50Mb | 16 | 25 | 29 | 30 | 100 |
|  | 200Mb | 5 | 10 | 41 | 44 | 100 |
|  | 1Gb | 0 | 4 | 3 | 3 | 10 |
| 1 per 5 kb | 1Mb | 69 | 7 | 9 | 15 | 100 |
|  | 10Mb | 49 | 8 | 18 | 25 | 100 |
|  | 50Mb | 10 | 4 | 35 | 51 | 100 |
|  | 200Mb | 0 | 0 | 48 | 52 | 100 |
|  | 1Gb | 0 | 0 | 5 | 5 | 10 |
| 1 per 50 kb | 1Mb | 83 | 7 | 4 | 6 | 100 |
|  | 10Mb | 83 | 5 | 4 | 8 | 100 |
|  | 50Mb | 59 | 10 | 10 | 21 | 100 |
|  | 200Mb | 20 | 6 | 40 | 34 | 100 |
|  | 1Gb | 0 | 0 | 4 | 6 | 10 |
| 1 per 100 kb | 1Mb | 86 | 4 | 3 | 7 | 100 |
|  | 10Mb | 78 | 5 | 7 | 10 | 100 |
|  | 50Mb | 64 | 6 | 14 | 16 | 100 |
|  | 200Mb | 43 | 6 | 30 | 21 | 100 |
|  | 1Gb | 0 | 1 | 5 | 4 | 10 |

**Supp Table 3:** Model selection in *fastsimcoal2* for 100 replicates, in which the true model was neutral equilibrium and selection was performed with 4 models: equilibrium, instantaneous size change, exponential size change, and instantaneous bottleneck. Varying densities of SNPs (all SNPs, 1 per 5 kb, 1 per 50 kb and 1 per 100 kb) were used to perform inference. Simulations were performed using msprime (0.7.3). Detailed methods including command lines can be found here: https://github.com/paruljohri/demographic_inference_with_selection/blob/main/CommandLines/SuppFigure4_SuppTable3_6.txt.

| Density of SNPs | Genome length | Equilibrium | Exponential change | Instantaneous change | Instantaneous Bottleneck | Total number of replicates |
| --- | --- | --- | --- | --- | --- | --- |
| all | 1Mb | 29 | 16 | 18 | 37 | 100 |
|  | 10Mb | 22 | 12 | 31 | 35 | 100 |
|  | 50Mb | 13 | 18 | 38 | 31 | 100 |
|  | 200Mb | 11 | 8 | 31 | 40 | 100 |
|  | 1Gb | 5 | 12 | 26 | 57 | 100 |
| 1 per 5 kb | 1Mb | 63 | 3 | 12 | 22 | 100 |
|  | 10Mb | 42 | 10 | 19 | 29 | 100 |
|  | 50Mb | 4 | 7 | 42 | 47 | 100 |
|  | 200Mb | 1 | 2 | 55 | 42 | 100 |
|  | 1Gb | 0 | 1 | 66 | 33 | 100 |
| 1 per 50 kb | 1Mb | 83 | 4 | 6 | 7 | 100 |
|  | 10Mb | 80 | 4 | 6 | 10 | 100 |
|  | 50Mb | 61 | 9 | 12 | 18 | 100 |
|  | 200Mb | 19 | 6 | 29 | 46 | 100 |
|  | 1Gb | 0 | 2 | 48 | 50 | 100 |
| 1 per 100 kb | 1Mb | 89 | 3 | 3 | 5 | 100 |
|  | 10Mb | 86 | 4 | 4 | 6 | 100 |
|  | 50Mb | 70 | 3 | 11 | 16 | 100 |
|  | 200Mb | 47 | 5 | 20 | 28 | 100 |
|  | 1Gb | 0 | 3 | 47 | 50 | 100 |

**Supp Table 4:** Linkage disequilibrium (*r*^!^) summarized in varying chromosomal sizes, and sampled in different densities, calculated using *Pylibseq* 0.2.3 in non-overlapping sliding windows across chromosomes with ∼10 SNPs per window. SNPs from separate chromosomes represent completely unlinked SNPs – 1 SNP was randomly sampled from each of 100 separate replicate chromosomes simulated, and this random sampling was performed 100 times. Detailed methods including command lines can be found here: https://github.com/paruljohri/demographic_inference_with_selection/blob/main/CommandLines/SuppTable4.txt.

| | | Number of SNPs per window | | $r^2$ | |
| --- | --- | --- | --- | --- | --- |
| Chromosome size | SNP density | mean | SD | mean | SD |
| **10Mb** | all | 10.7 | 3.6 | 0.1181 | 0.0885 |
|  | 1 per 5 kb | 8.3 | 0.7 | 0.0731 | 0.0504 |
|  | 1 per 50 kb | 9.8 | 0.4 | 0.0271 | 0.0141 |
|  | 1 per 100 kb | 10.0 | 0.2 | 0.0203 | 0.0101 |
|  | separate chromosomes | 10.5 | 0.5 | 0.0103 | 0.0047 |
| **1 Mb** | all | 10.7 | 3.6 | 0.1174 | 0.0889 |
|  | 1 per 5 kb | 8.4 | 0.7 | 0.0725 | 0.0503 |
|  | 1 per 50 kb | 10.0 | 0.0 | 0.0252 | 0.0108 |
|  | 1 per 100 kb | 10.0 | 0.0 | 0.0193 | 0.0080 |
|  | separate chromosomes | 10.5 | 0.5 | 0.0102 | 0.0047 |
| **50Mb** | all | 10.7 | 3.6 | 0.1181 | 0.0888 |
|  | 1 per 5 kb | 8.3 | 0.7 | 0.0729 | 0.0502 |
|  | 1 per 50 kb | 9.8 | 0.4 | 0.0269 | 0.0136 |
|  | 1 per 100 kb | 9.9 | 0.3 | 0.0203 | 0.0099 |
|  | separate chromosomes | 10.5 | 0.5 | 0.0101 | 0.0046 |
| **200 Mb** | all | 10.6 | 3.6 | 0.1180 | 0.0888 |
|  | 1 per 5 kb | 8.3 | 0.7 | 0.0728 | 0.0499 |
|  | 1 per 50 kb | 9.8 | 0.4 | 0.0269 | 0.0137 |
|  | 1 per 100 kb | 9.9 | 0.3 | 0.0202 | 0.0098 |
|  | separate chromosomes | 10.5 | 0.5 | 0.0101 | 0.0046 |
| **1 Gb** | all | 10.6 | 3.6 | 0.1181 | 0.0891 |
|  | 1 per 5 kb | 8.4 | 0.7 | 0.0727 | 0.0495 |
|  | 1 per 50 kb | 9.8 | 0.4 | 0.0269 | 0.0134 |
|  | 1 per 100 kb | 9.9 | 0.3 | 0.0204 | 0.0102 |
|  | separate chromosomes | 6.5 | 0.5 | 0.0092 | 0.0046 |

**Supp Table 5:**
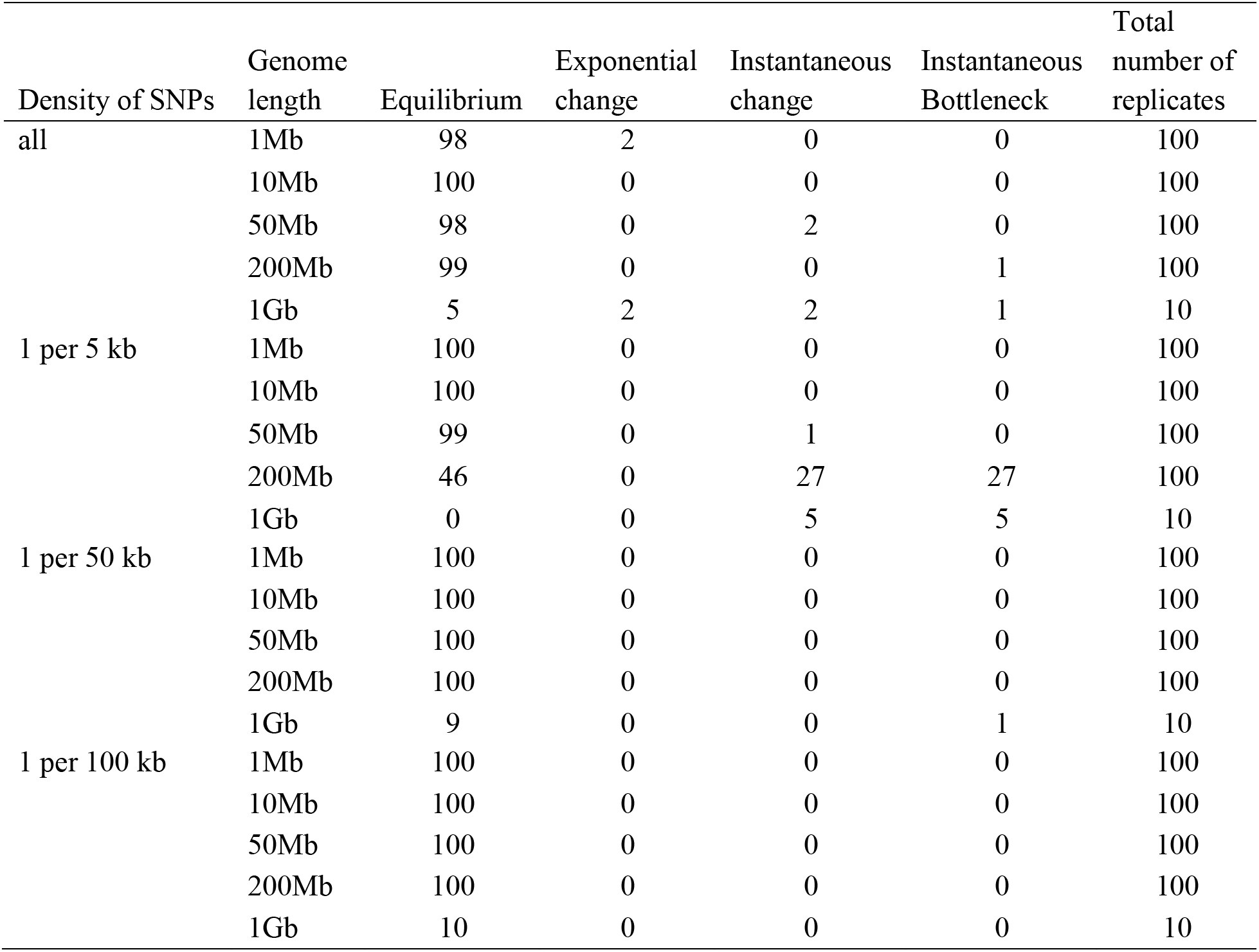
Model selection with a higher AIC penalty (=25×the number of parameters) in which the true model was neutral equilibrium, and model selection was performed with 4 models: equilibrium, instantaneous size change, exponential size change, and instantaneous bottleneck. Varying densities of SNPs (all SNPs, 1 per 5 kb, 1 per 50 kb and 1 per 100 kb) were used to perform inference. Simulations were performed in SLiM (3.1) for 10*N* generations. Detailed methods including command lines can be found here: https://github.com/paruljohri/demographic_inference_with_selection/blob/main/CommandLines/SuppFigure3_SuppTable2_5.txt.

| Density of SNPs | Genome length | Equilibrium | Exponential change | Instantaneous change | Instantaneous Bottleneck | Total number of replicates |
| --- | --- | --- | --- | --- | --- | --- |
| all | 1Mb | 98 | 2 | 0 | 0 | 100 |
|  | 10Mb | 100 | 0 | 0 | 0 | 100 |
|  | 50Mb | 98 | 0 | 2 | 0 | 100 |
|  | 200Mb | 99 | 0 | 0 | 1 | 100 |
|  | 1Gb | 5 | 2 | 2 | 1 | 10 |
| 1 per 5 kb | 1Mb | 100 | 0 | 0 | 0 | 100 |
|  | 10Mb | 100 | 0 | 0 | 0 | 100 |
|  | 50Mb | 99 | 0 | 1 | 0 | 100 |
|  | 200Mb | 46 | 0 | 27 | 27 | 100 |
|  | 1Gb | 0 | 0 | 5 | 5 | 10 |
| 1 per 50 kb | 1Mb | 100 | 0 | 0 | 0 | 100 |
|  | 10Mb | 100 | 0 | 0 | 0 | 100 |
|  | 50Mb | 100 | 0 | 0 | 0 | 100 |
|  | 200Mb | 100 | 0 | 0 | 0 | 100 |
|  | 1Gb | 9 | 0 | 0 | 1 | 10 |
| 1 per 100 kb | 1Mb | 100 | 0 | 0 | 0 | 100 |
|  | 10Mb | 100 | 0 | 0 | 0 | 100 |
|  | 50Mb | 100 | 0 | 0 | 0 | 100 |
|  | 200Mb | 100 | 0 | 0 | 0 | 100 |
|  | 1Gb | 10 | 0 | 0 | 0 | 10 |

**Supp Table 6:** Model selection with a higher AIC penalty (=25×the number of parameters) in which the true model was neutral equilibrium, and model selection was performed with 4 models: equilibrium, instantaneous size change, exponential size change, and instantaneous bottleneck. Varying densities of SNPs (all SNPs, 1 per 5 kb, 1 per 50 kb and 1 per 100 kb) were used to perform inference. Simulations were performed using msprime (0.7.3). Detailed methods including command lines can be found here: https://github.com/paruljohri/demographic_inference_with_selection/blob/main/CommandLines/SuppFigure4_SuppTable3_6.txt.

| Density of SNPs | Genome length | Equilibrium | Exponential change | Instantaneous change | Instantaneous Bottleneck | Total number of replicates |
| --- | --- | --- | --- | --- | --- | --- |
| all | 1Mb | 100 | 0 | 0 | 0 | 100 |
|  | 10Mb | 100 | 0 | 0 | 0 | 100 |
|  | 50Mb | 97 | 0 | 2 | 1 | 100 |
|  | 200Mb | 96 | 0 | 4 | 0 | 100 |
|  | 1Gb | 93 | 1 | 3 | 3 | 100 |
| 1 per 5 kb | 1Mb | 100 | 0 | 0 | 0 | 100 |
|  | 10Mb | 100 | 0 | 0 | 0 | 100 |
|  | 50Mb | 97 | 1 | 0 | 2 | 100 |
|  | 200Mb | 45 | 1 | 30 | 24 | 100 |
|  | 1Gb | 0 | 1 | 66 | 33 | 100 |
| 1 per 50 kb | 1Mb | 100 | 0 | 0 | 0 | 100 |
|  | 10Mb | 100 | 0 | 0 | 0 | 100 |
|  | 50Mb | 100 | 0 | 0 | 0 | 100 |
|  | 200Mb | 100 | 0 | 0 | 0 | 100 |
|  | 1Gb | 89 | 1 | 4 | 6 | 100 |
| 1 per 100 kb | 1Mb | 100 | 0 | 0 | 0 | 100 |
|  | 10Mb | 100 | 0 | 0 | 0 | 100 |
|  | 50Mb | 100 | 0 | 0 | 0 | 100 |
|  | 200Mb | 100 | 0 | 0 | 0 | 100 |
|  | 1Gb | 100 | 0 | 0 | 0 | 100 |

**Supp Table 7:** Parameters underlying the human-like demographic models considered.

| | **Demographic models** | **Ancestral population size**<br>($N_{anc}$) | **Current population size**<br>($N_{cur}$) | **Time of change in generations** |
| --- | --- | --- | --- | --- |
| 1 | Equilibrium | 10,000 | 10,000 | NA |
| 2 | Exponential growth | 1000 | 30,000 | 850 |
| 3 | Instantaneous decline | 12,300 | 2,100 | 4,750 |

**Supp Table 8:** Comparison of inference of parameters using *fastsimcoal2* under neutrality when inference is performed using all SNPs *vs*. using sparser SNP sampling (1 per 5 kb). In this example, 20% of the genome is exonic, and these exonic regions were masked when performing inference.

| **Demographic model** | **Parameter** | **True value** | **Inferred value using all SNPs**<br>(Mean±SD) | **Inferred value using 1 SNP per 5 kb**<br>(Mean±SD) |
| --- | --- | --- | --- | --- |
| Equilibrium | $N_{anc}$ | 10,000 | 9,781 ± 23 | 9,378 ± 567 |
| Exponential growth | $N_{anc}$ | 1000 | 977 ± 3 | 963 ± 4 |
| | $N_{cur}$ | 30,000 | 25,489 ± 123 | 25,641 ± 184 |
| | Time of change | 850 | 808 ± 3 | 816 ± 4 |
| Instantaneous decline | $N_{anc}$ | 12,300 | 11,978 ± 463 | 11,513 ± 894 |
| | $N_{cur}$ | 2,100 | 2,097 ± 15 | 2,681 ± 12 |
| | Time of change | 4,750 | 4,703 ± 246 | 7,458 ± 596 |

**Supp Table 9:** Nucleotide diversity in the presence of BGS relative to that under neutrality (*B*), calculated for a neutral site distance *y* bases from the end of a gene/exon of length 500 bp. The exon experiences purifying selection with strength 2*N*_e_*s*=10, where *N*_e_ is the effective population size and *s* is the reduction in fitness. Shown below is *t*=*hs* where *h* is the dominance coefficient (assumed to be 0.5 here), *r* is the recombination rate per site per generation, and *u* is the mutation rate per site per generation. *B* was calculated using Equation 2 of Johri *et al*. (2020).

| $N_e$ | $t(2N_e s$<br>$=10)$ | $r$ | $u$ | $B(y=1)$ | $B(y=10)$ | $B$<br>$(y=1000)$ |
| --- | --- | --- | --- | --- | --- | --- |
| $10^4$ | 0.00025 | $1.00 \times 10^{-8}$ | $1.00 \times 10^{-8}$ | 0.9805 | 0.9806 | 0.9820 |
| | | $1.00 \times 10^{-6}$ | $1.00 \times 10^{-6}$ | 0.5152 | 0.5312 | 0.9444 |
| $10^6$ | $2.5 \times 10^{-6}$ | $1.00 \times 10^{-8}$ | $1.00 \times 10^{-8}$ | 0.5152 | 0.5312 | 0.9445 |
| | | $1.00 \times 10^{-6}$ | $1.00 \times 10^{-6}$ | 0.4920 | 0.8227 | 0.9992 |

**Supp Figure 1:**
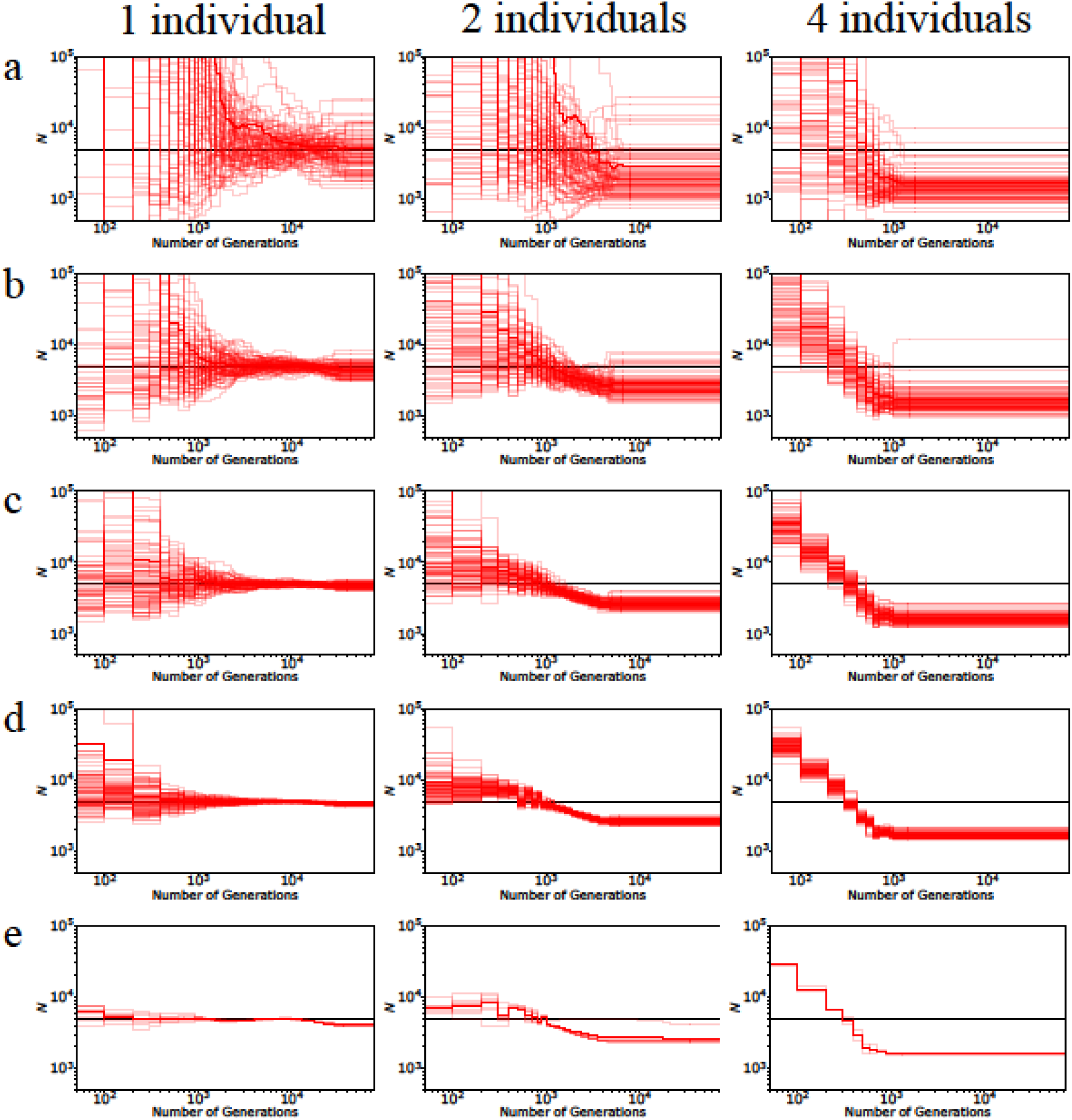
Performance of MSMC under neutrality and demographic equilibrium when using 1, 2, and 4 diploid individuals for inference, for varying chromosome sizes: (a) 1Mb, (b) 10 Mb, (c) 50 Mb, (d) 200 Mb, (e) 1 Gb. Simulations were performed using SLiM (3.1) for 10*N* generations with 100 replicates for panels (a-d) and 10 for panel (e). Detailed methods including command lines can be found here: https://github.com/paruljohri/demographic_inference_with_selection/blob/main/CommandLines/SuppFigure1.txt.

**Supp Figure 2:**
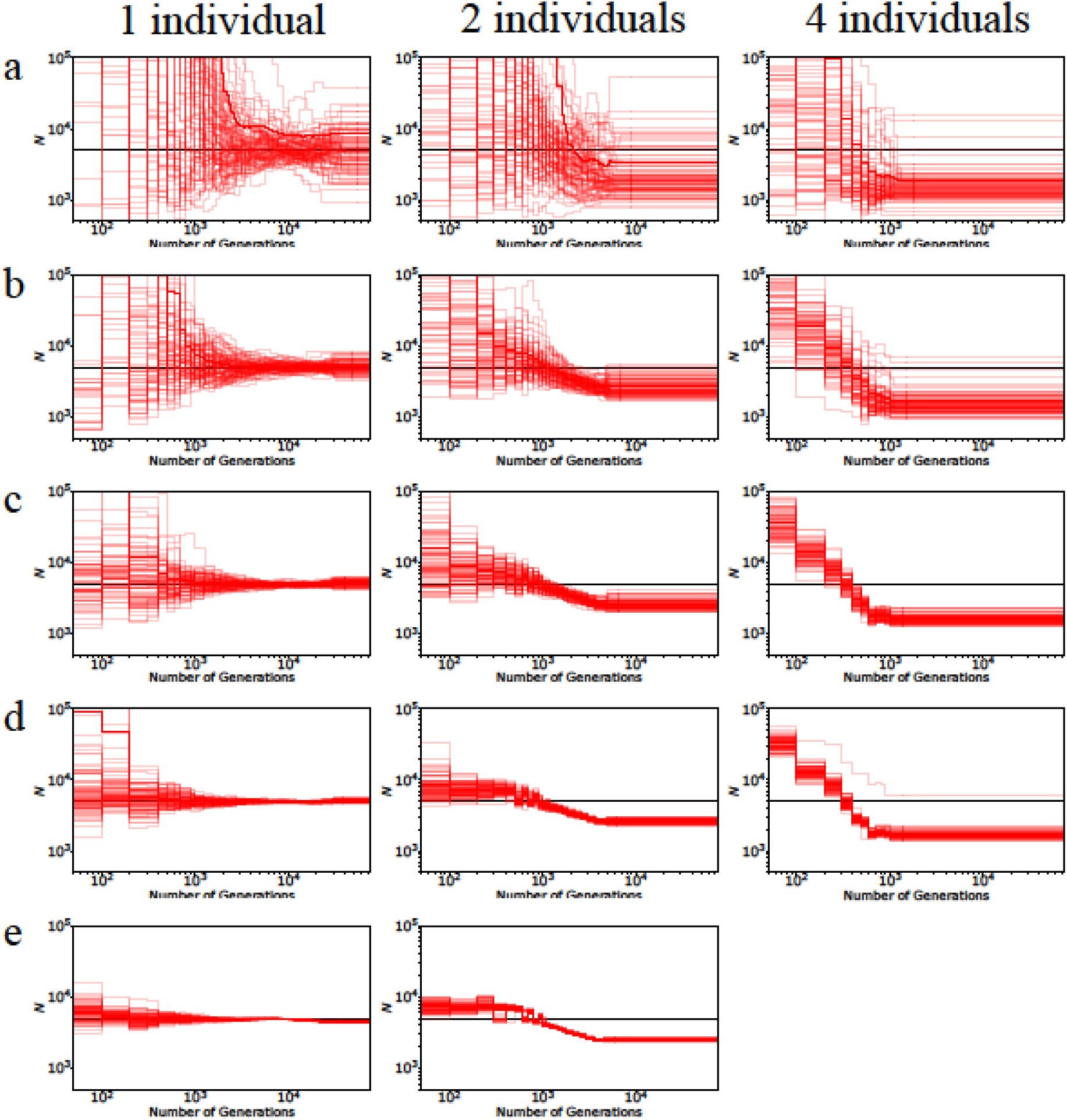
Performance of MSMC under neutrality and demographic equilibrium when using 1, 2, and 4 diploid individuals for inference, for varying chromosome sizes: (a) 1 Mb, (b) 10 Mb, (c) 50 Mb, (d) 200 Mb, (e) 1 Gb. Simulations were performed using msprime (0.7.3) with 100 replicates for each panel. MSMC runs with 4 diploid individuals at 1Gb could not be obtained due to the long computational times required. Detailed methods including command lines can be found here: https://github.com/paruljohri/demographic_inference_with_selection/blob/main/CommandLines/SuppFigure2.txt.

**Supp Figure 3:**
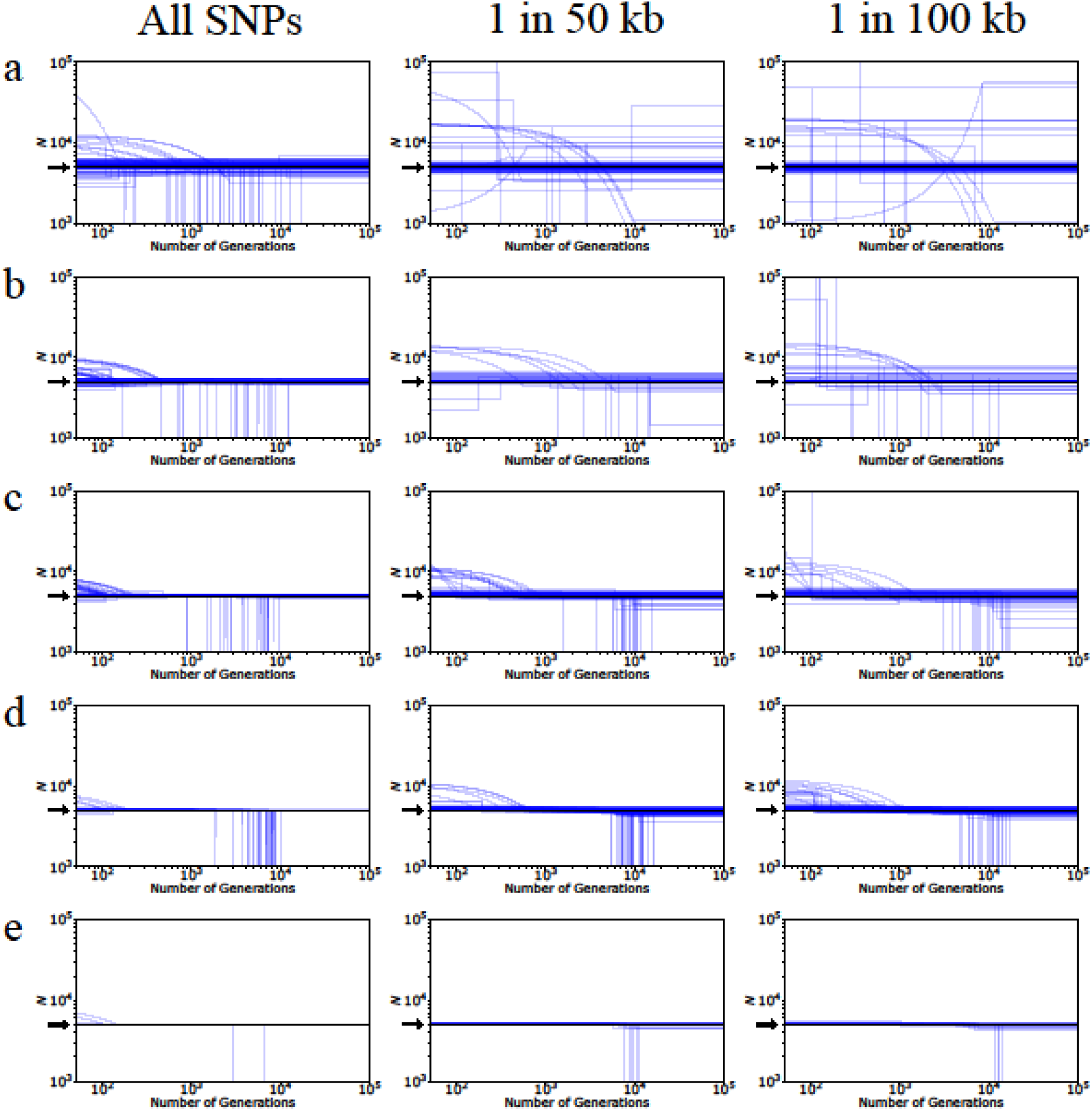
Performance of *fastsimcoal2* under neutrality and demographic equilibrium when simulations were performed using SLiM (3.1) for varying chromosome sizes: (a) 1 Mb, (b) 10 Mb, (c) 50 Mb, (d) 200 Mb, (e) 1 Gb. Model selection was performed with 4 models – equilibrium, instantaneous size change, exponential size change, and instantaneous bottleneck. Inference was performed using 50 diploid individuals using all SNPs (left column), 1 SNP per 50 kb (middle column) and 1 SNP per 100 kb (right column). The inferred population size estimates of the best model are plotted (blue lines). The numbers of replicates for (a)-(d) were 100, while for (e) were 10. The true model is shown in black and indicated by the position of the arrow. Detailed methods including command lines can be found here: https://github.com/paruljohri/demographic_inference_with_selection/blob/main/CommandLines/SuppFigure3_SuppTable2_5.txt.

**Supp Figure 4:**
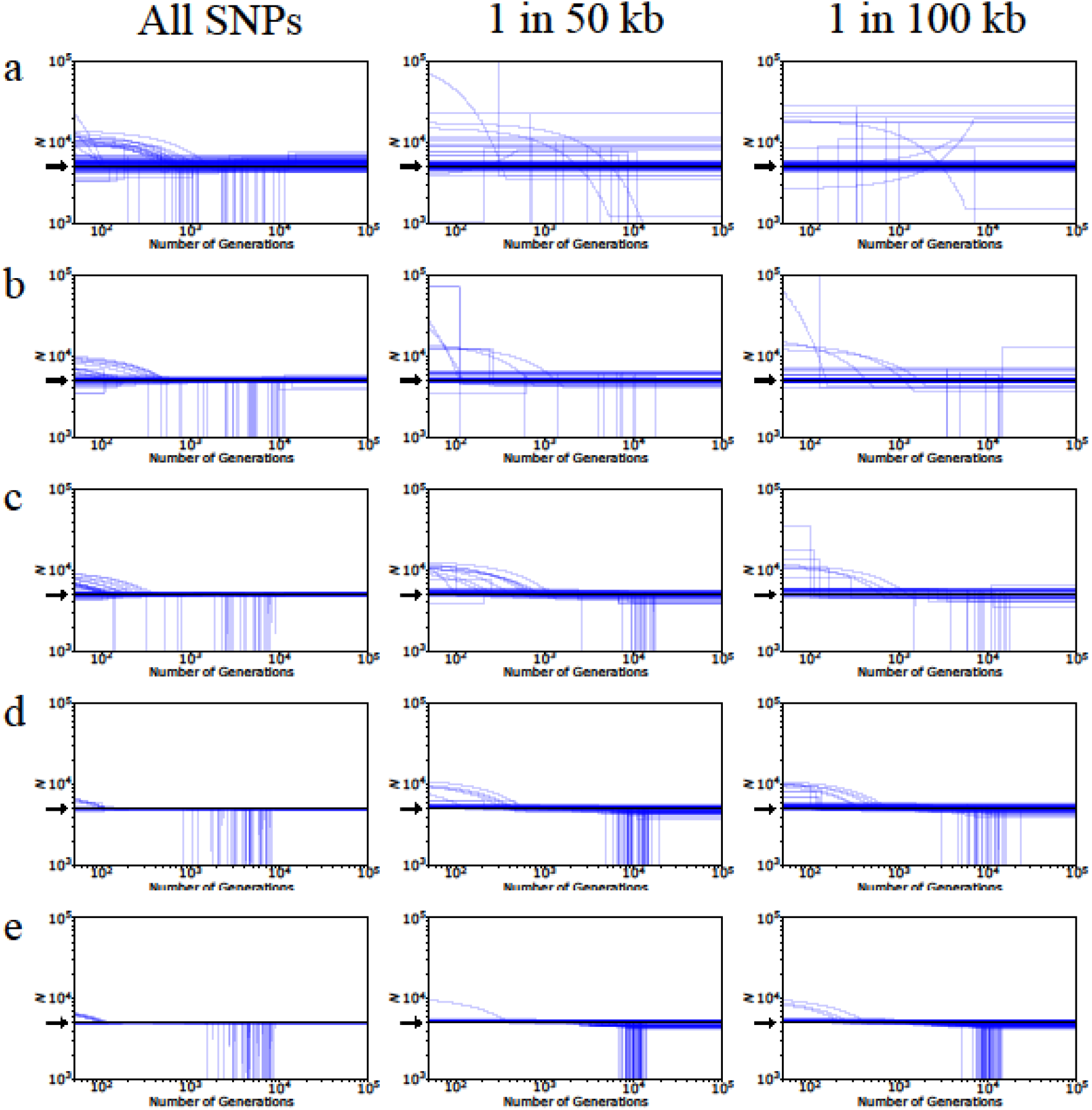
Performance of *fastsimcoal2* under neutrality and demographic equilibrium when simulations were performed using msprime (0.7.3) for varying chromosome sizes: (a) 1 Mb, (b) 10 Mb, (c) 50 Mb, (d) 200 Mb, (e) 1 Gb. Model selection was performed with 4 models – equilibrium, instantaneous size change, exponential size change, and instantaneous bottleneck. Inference was performed using 50 diploid individuals using all SNPs (left column), 1 SNP per 50 kb (middle column) and 1 SNP per 100 kb (right column). The inferred population size estimates of the best model are plotted (blue lines), with 100 replicates for each panel. The true model is shown in black and indicated by the position of the arrow. Detailed methods including command lines can be found here: https://github.com/paruljohri/demographic_inference_with_selection/blob/main/CommandLines/SuppFigure4_SuppTable3_6.txt.

**Supp Figure 5:**
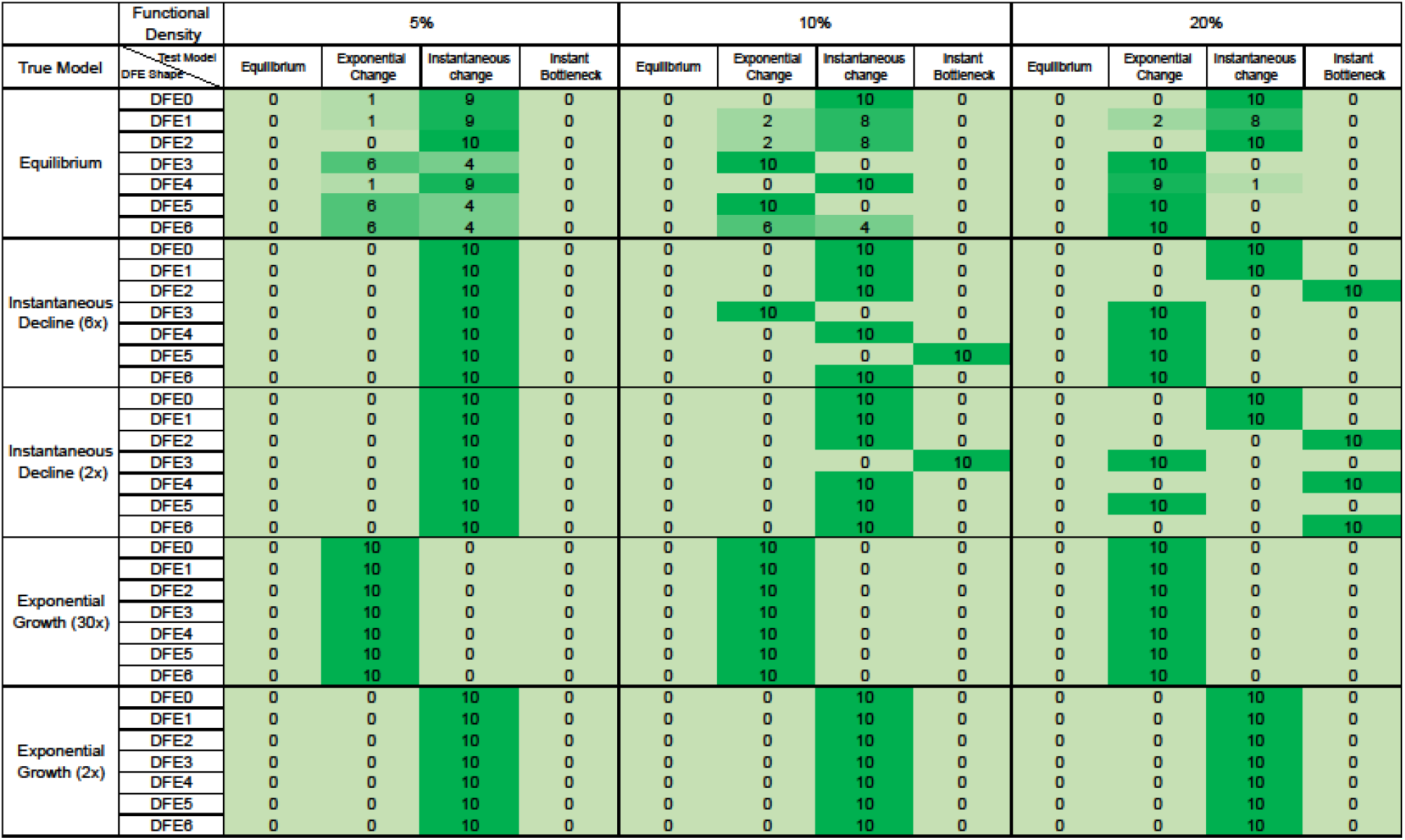
Model selection by *fastsimcoal2* in the presence of BGS, when chosen from four possible models: equilibrium, instantaneous size change, exponential size change, and instantaneous bottleneck. The DFEs are specified in Table 1, and DFE0 refers to neutrality. Results are shown when all SNPs were used for inference, when 5%, 10% or 20% of the genomes were exonic (with exonic sites masked), and the standard AIC penalty was used. Detailed methods including command lines can be found here: https://github.com/paruljohri/demographic_inference_with_selection/blob/main/CommandLines/SuppFigure5.txt.

**Supp Figure 6:**
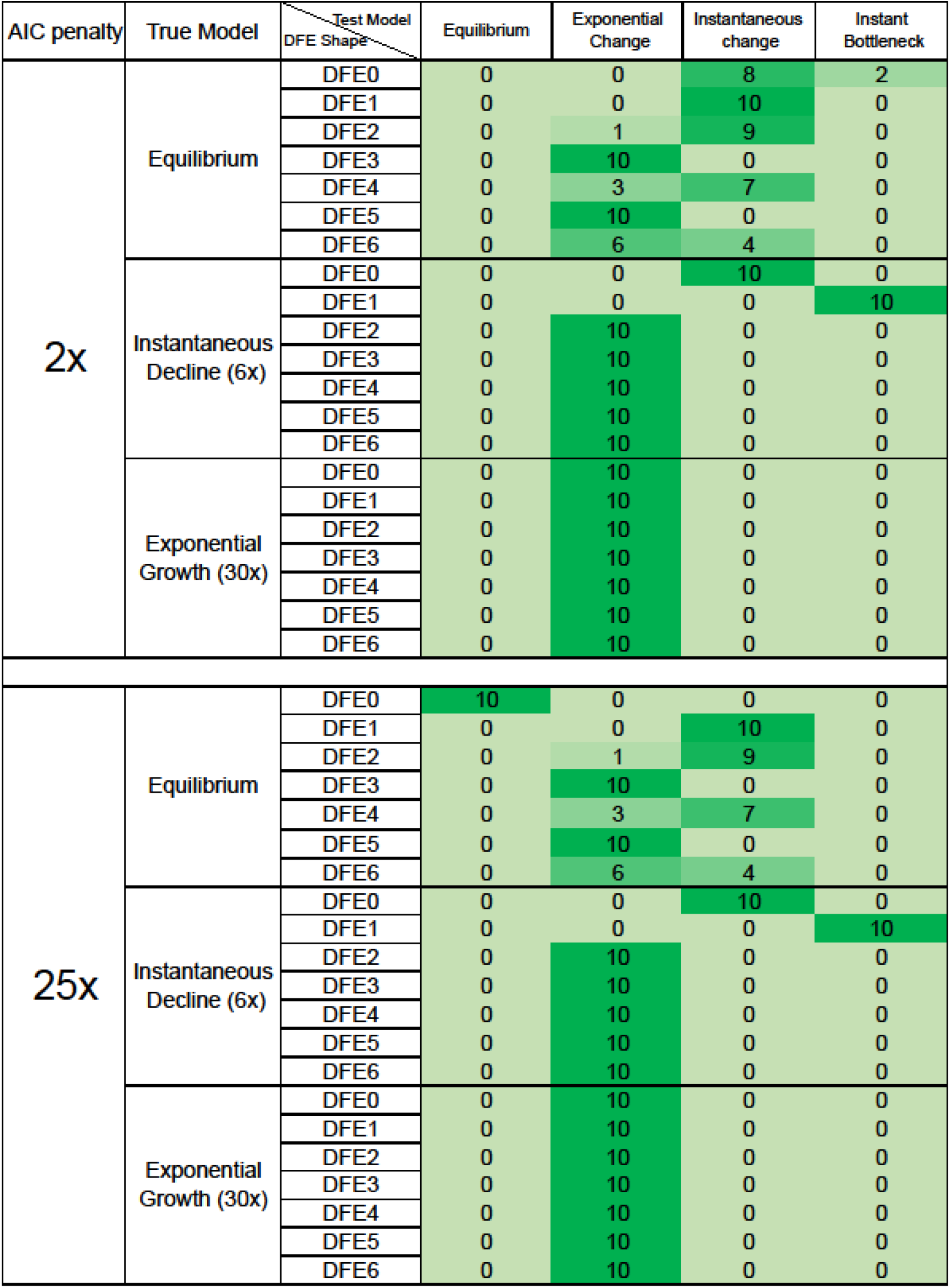
Model selection by *fastsimcoal2* in the presence of BGS, when chosen from four possible models: equilibrium, instantaneous size change, exponential size change, and instantaneous bottleneck. The DFEs are specified in Table 1, and DFE0 refers to neutrality. Results are shown when SNPs used for inference were separated at a distance of 100 kb, 20% of the genomes were exonic, and the AIC penalty was 2× (standard) or 25× the number of parameters. Detailed methods including command lines can be found here: https://github.com/paruljohri/demographic_inference_with_selection/blob/main/CommandLines/SuppFigure6.txt.

**Supp Figure 7:**
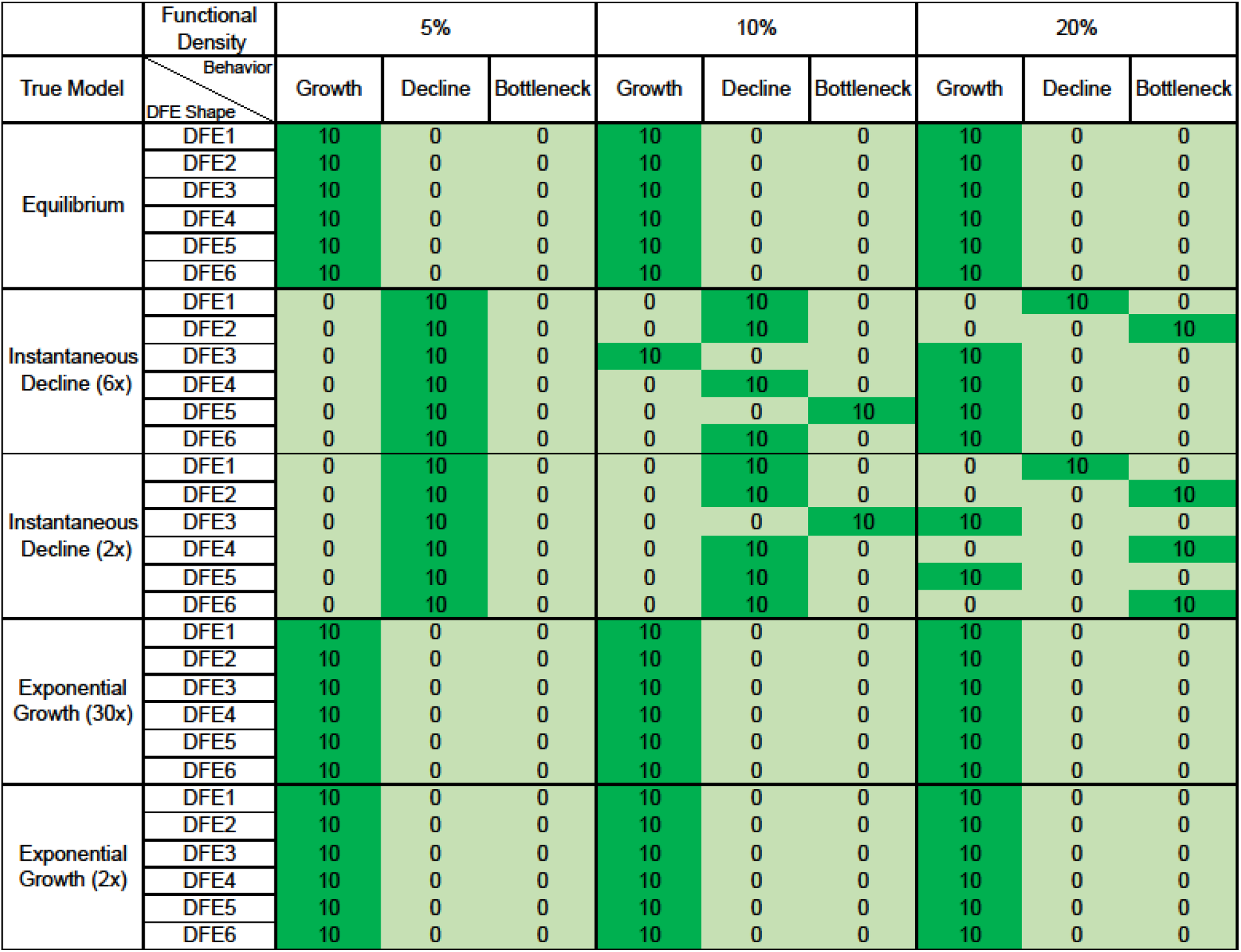
Effects of BGS on inference of growth or decline by *fastsimcoal2*. The inferred model was classified as growth if *N_anc_* < *N_cur_* and as decline if *N_anc_* > *N_cur_*. The DFEs are specified in Table 1. Results are shown for all SNPs, when 5%, 10% and 20% of the genomes were exonic (with exonic sites masked), and a standard AIC penalty was used. Detailed methods including command lines can be found here: https://github.com/paruljohri/demographic_inference_with_selection/blob/main/CommandLines/SuppFigure7.txt.

**Supp Figure 8:**
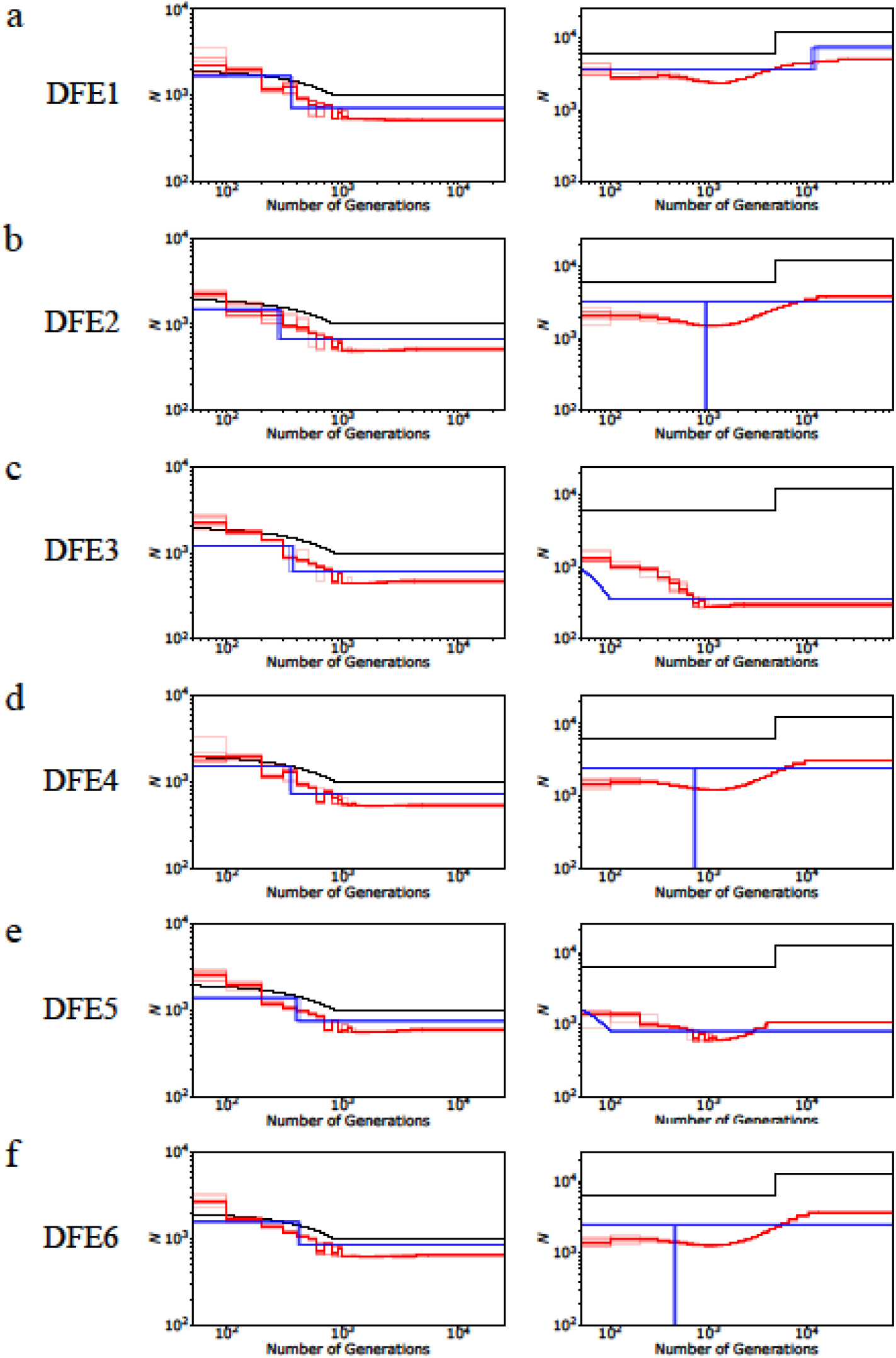
Inferred demography from MSMC (red lines) and *fastsimcoal2* (blue lines) in the presence of background selection, with the true DFE shown to the left of the panel, for 2-fold instantaneous decline (right column) and 2-fold exponential growth (left column). In this case, 20% of the genome was exonic (and exonic sites were masked). The true demographic models are shown as black lines. Detailed methods including command lines can be found here: https://github.com/paruljohri/demographic_inference_with_selection/blob/main/CommandLines/SuppFigure8.txt.

**Supp Figure 9:**
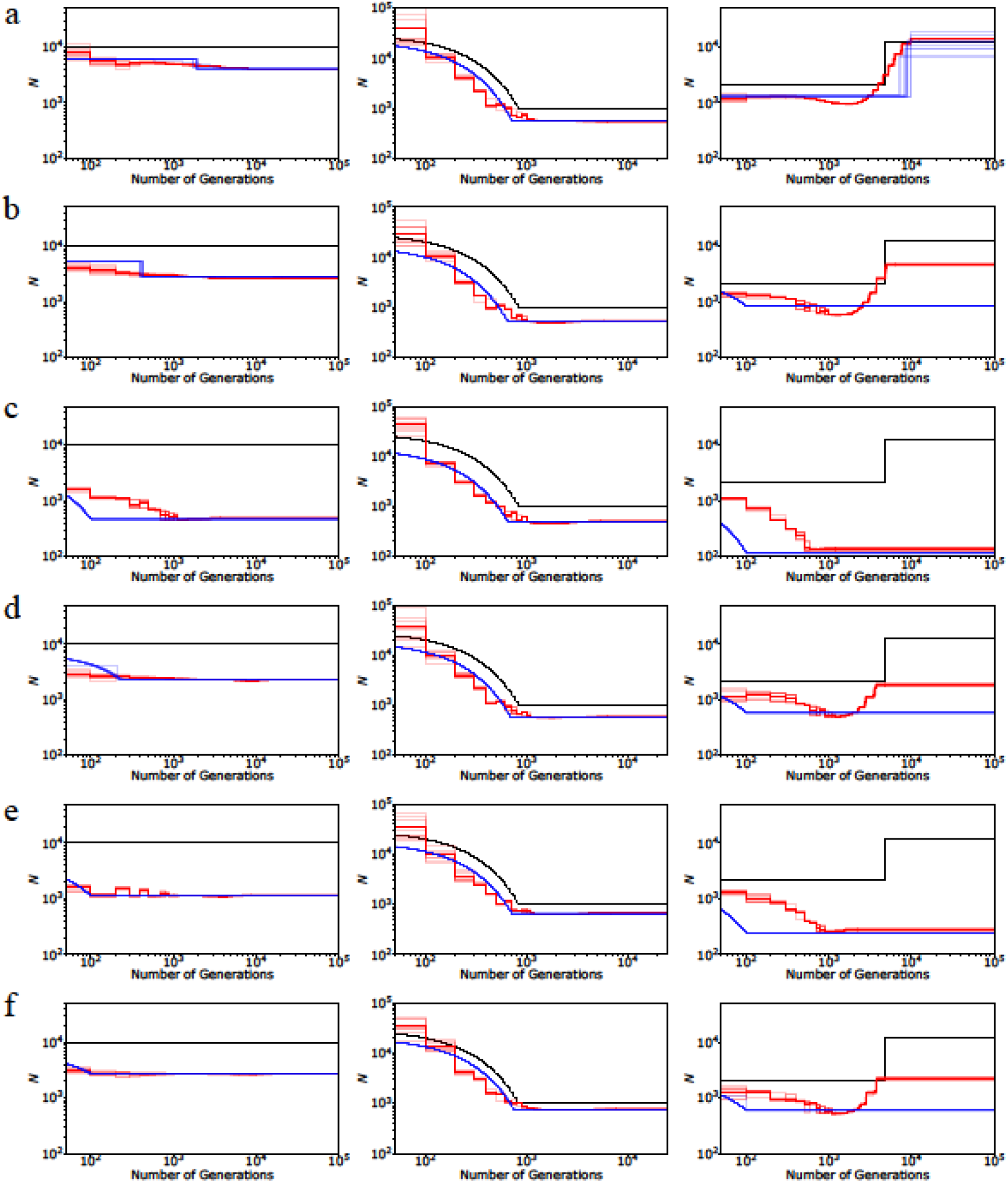
Inference of demography by MSMC (red lines; 10 replicates) and *fastsimcoal2* (blue lines; 10 replicates) under demographic equilibrium (left column), 30-fold exponential growth (middle column), and ∼6-fold instantaneous decline (right column) in the presence of direct purifying selection (*i.e*., directly selected sites are not masked). The true demographic model is depicted in black lines. Exonic sites experience purifying selection specified by the following DFEs (defined in Table 1): (a) DFE1, (b) DFE2, (c) DFE3, (d) DFE4, (e) DFE5, (f) DFE6. In this case, 20% of the genome was exonic and all SNPs were used for inference, including exonic sites. Detailed methods including command lines can be found here: https://github.com/paruljohri/demographic_inference_with_selection/blob/main/CommandLines/SuppFigure9.txt.

**Supp Figure 10:**
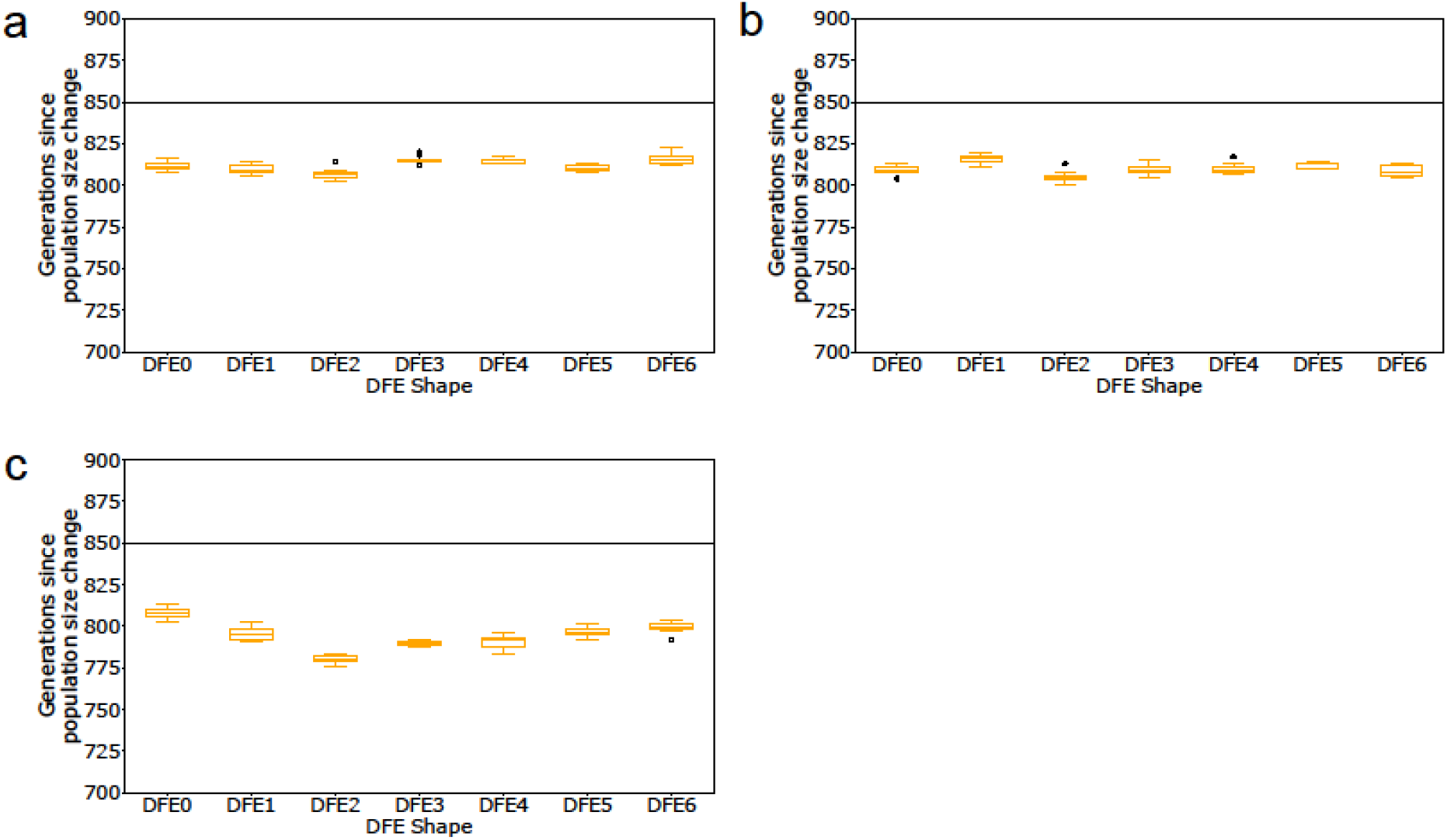
Inference of the timing of 30-fold exponential growth by *fastsimcoal2* in the presence of BGS compared to neutrality (DFE0). The DFEs are specified in Table 1. Results are shown for the case where (a) 5%, (b) 10%, and (c) 20% of the genome is exonic (with exonic sites masked), and all non-exonic SNPs are used for inference. The black horizontal line represents the true timing.

**Supp Figure 11:**
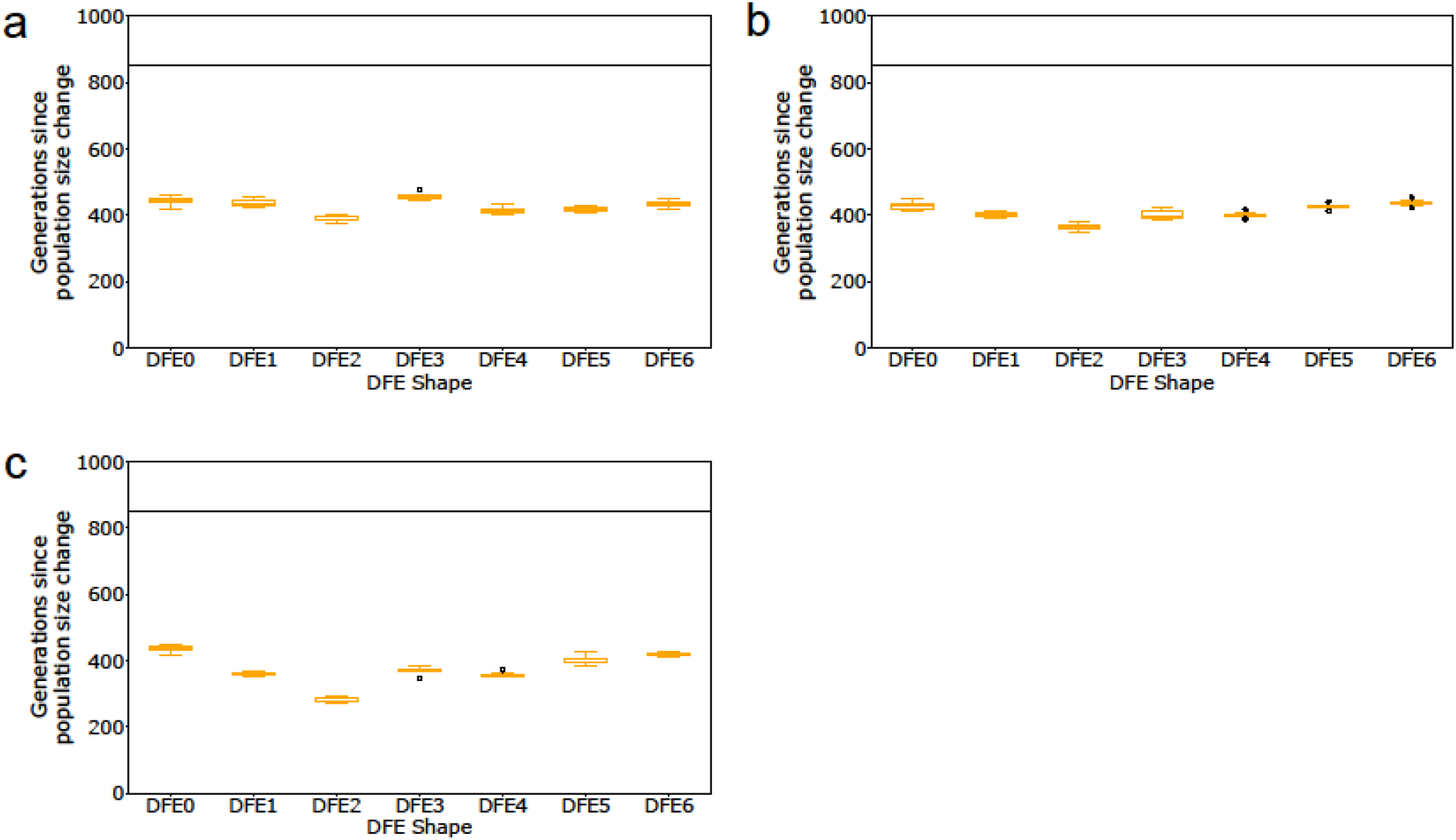
Inference of the timing of 2-fold exponential growth by *fastsimcoal2* in the presence of BGS compared to neutrality (DFE0). Results are shown for the case where (a) 5%, (b) 10%, and (c) 20% of the genome is exonic (with exonic sites masked), and all non-exonic SNPs are used for inference. The black horizontal line represents the true timing.

**Supp Figure 12:**
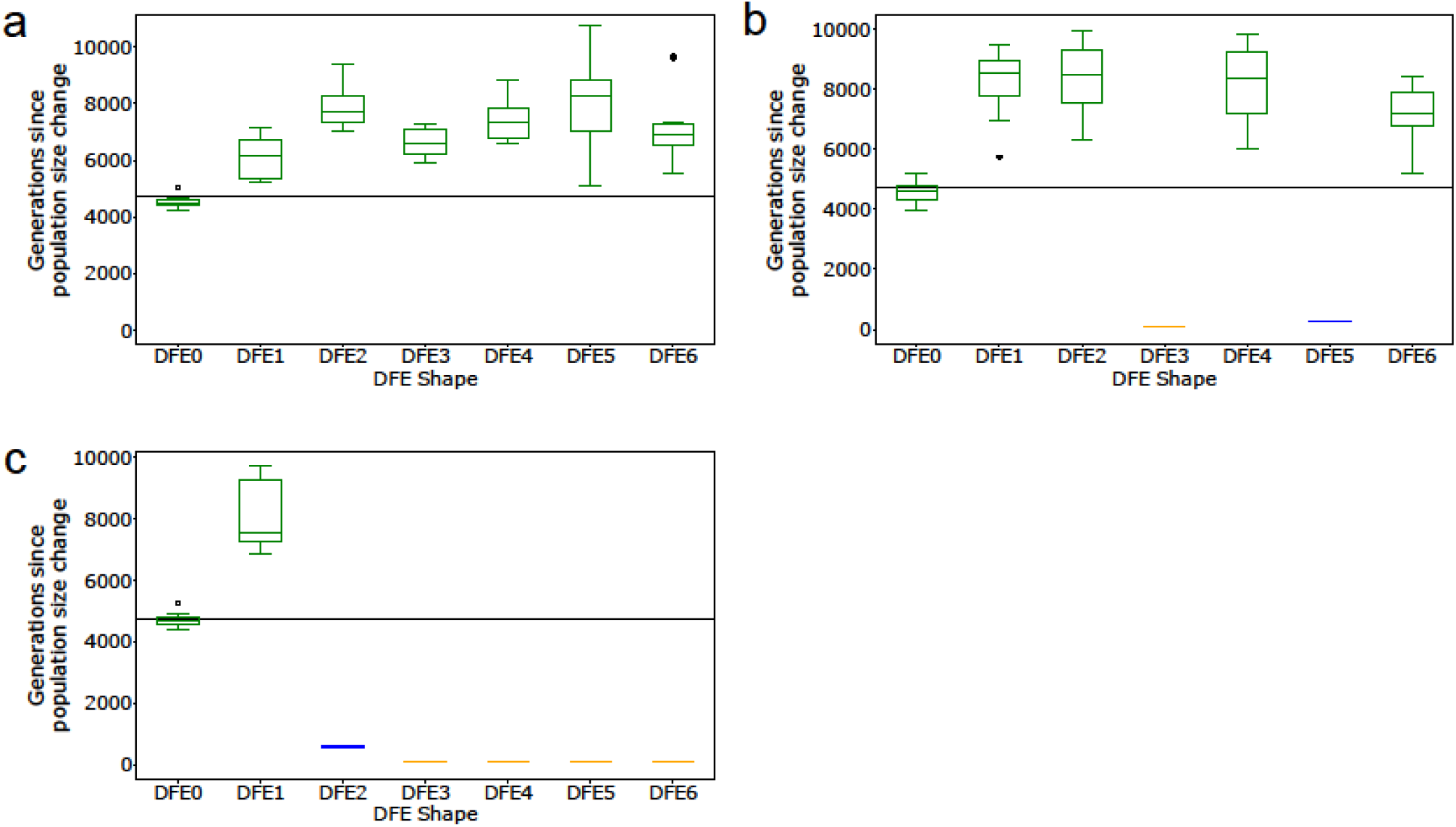
Inference of the timing of 6-fold instantaneous decline by *fastsimcoal2* in the presence of BGS compared to neutrality (DFE0). The DFEs are specified in Table 1. Results are shown for the case in which (a) 5%, (b) 10%, and (c) 20% of the genome is exonic (with exonic sites masked), and all non-exonic SNPs are used for inference. The black horizontal line represents the true timing. Boxplots are presented in green if decline was inferred, in yellow if growth was inferred, and in blue if a bottleneck was inferred.

**Supp Figure 13:**
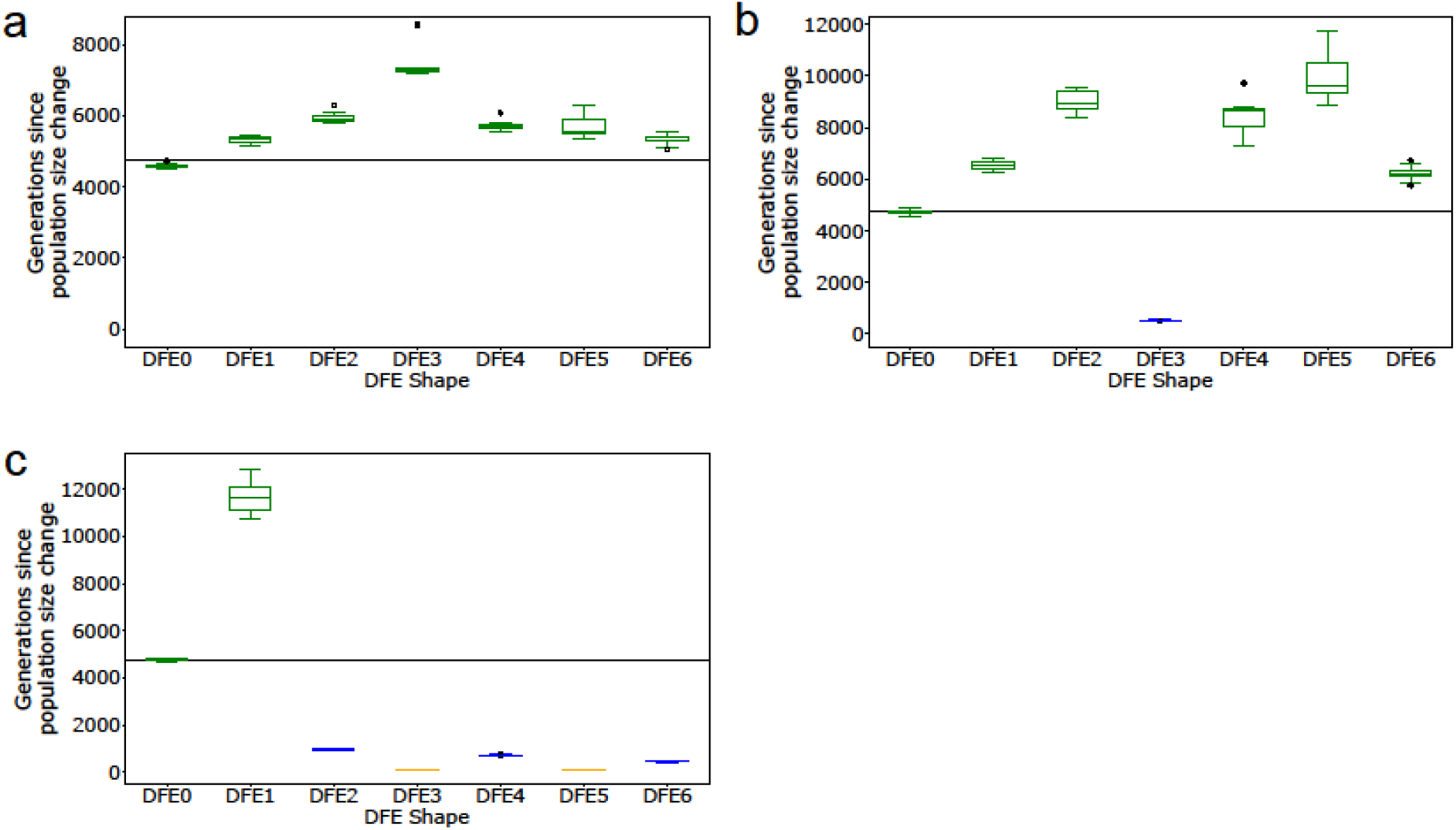
Inference of the timing of 2-fold instantaneous decline by *fastsimcoal2* in the presence of BGS compared to neutrality (DFE0). The DFEs are specified in Table 1. Results are shown for the case where (a) 5%, (b) 10%, and (c) 20% of the genome is exonic (with exonic sites masked), and all non-exonic SNPs are used for inference. The black horizontal line represents the true timing. Boxplots are presented in green if decline was inferred, in yellow if growth was inferred, and in blue if a bottleneck was inferred.

**Supp Figure 14:**
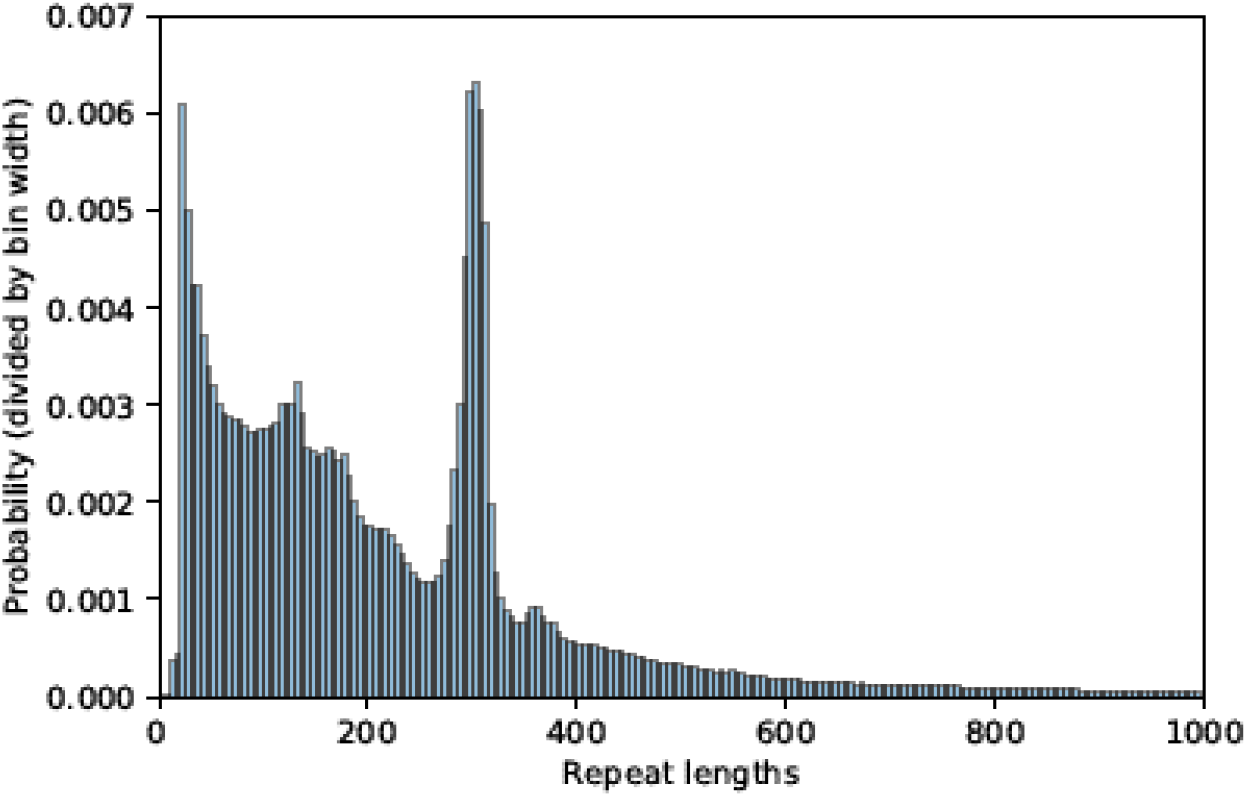
Distribution of lengths of repeat regions in the human genomes (*hg19*). Shown above is the distribution of lengths up to 1000 bp, although lengths of repeat regions range between 6 to 160602 bp. Detailed methods including command lines can be found here: https://github.com/paruljohri/demographic_inference_with_selection/blob/main/CommandLines/SuppFigure14.txt.

**Supp Figure 15:**
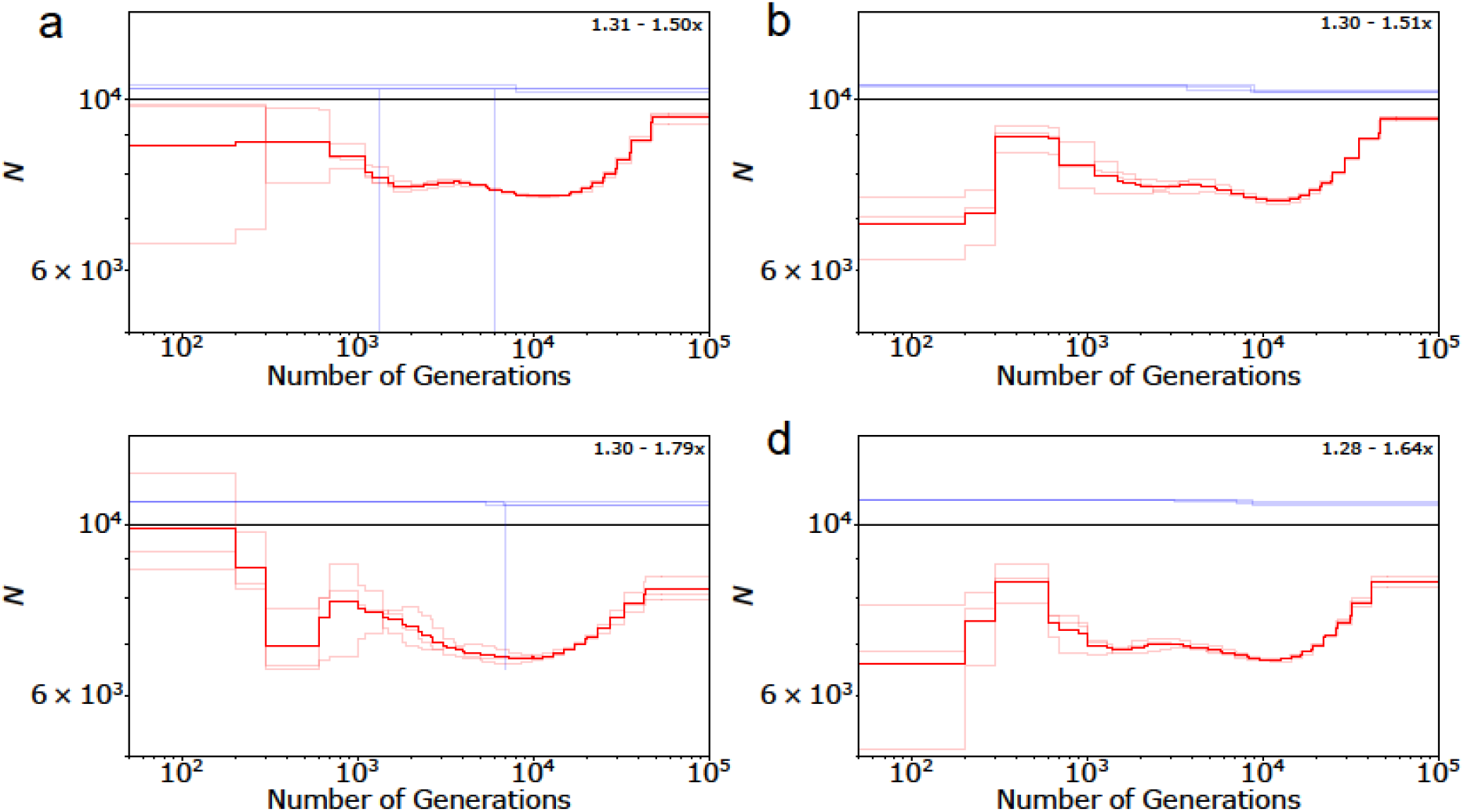
Performance of demographic inference by MSMC (red lines) and *fastsimcoal2* (blue lines) under different scenarios of neutrality, when the true model is equilibrium: (a) there is variation in recombination and mutation rates, (b) there is variation in recombination and mutation rates and the centromeric region is masked, (c) there is variation in recombination and mutation rates, and short regions resembling repeats (comprising 10% of each chromosome) are randomly masked across the genome, and (d) there is variation in recombination and mutation rates, and the centromere as well as small-sized repeats are randomly masked across the genome. The maximum and minimum fold change detected in every scenario is indicated on the upper right corner. Detailed methods including command lines can be found here: https://github.com/paruljohri/demographic_inference_with_selection/blob/main/CommandLines/SuppFigure15-20.txt.

**Supp Figure 16:**
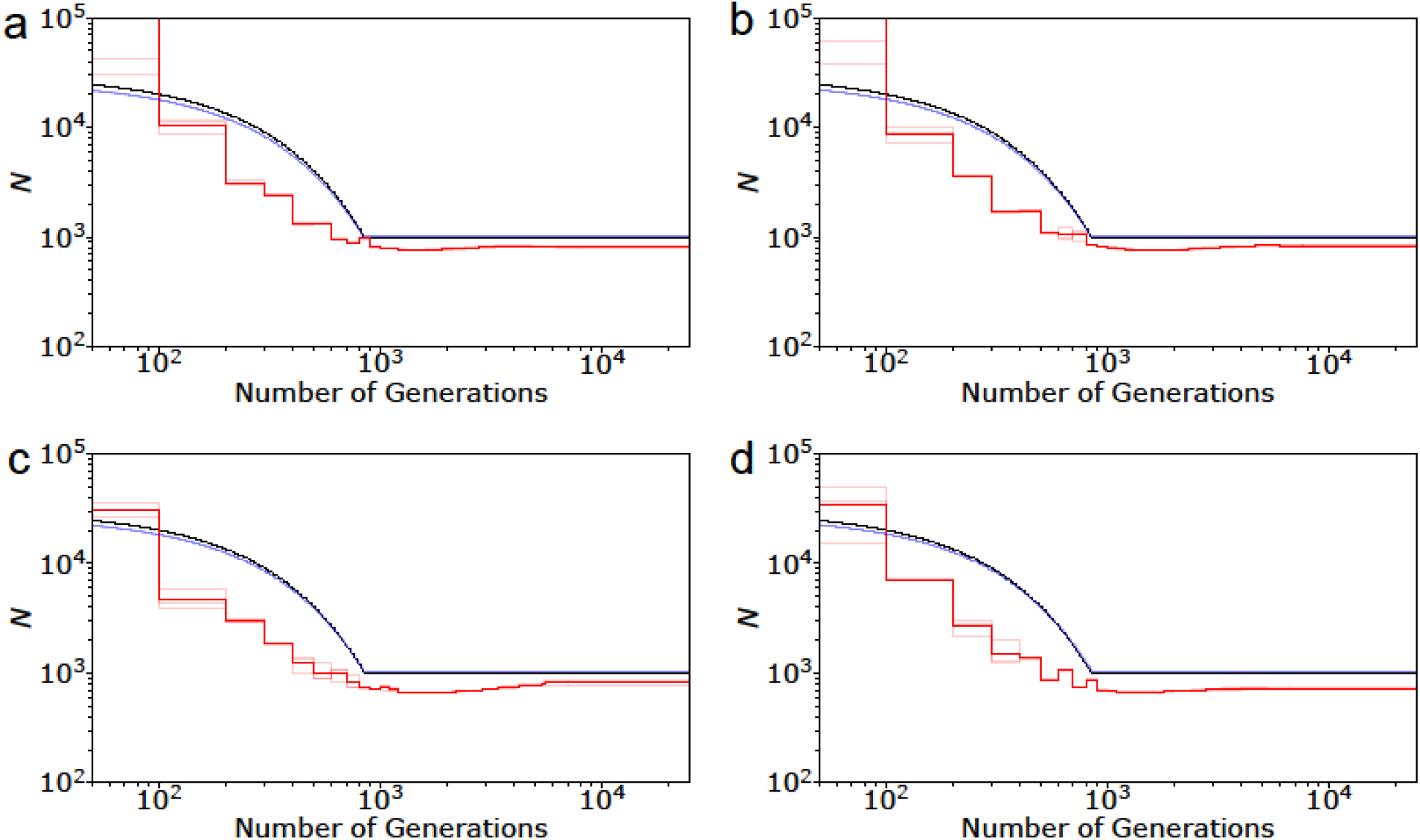
Performance of demographic inference by MSMC (red lines) and *fastsimcoal2* (blue lines) under different scenarios of neutrality, when the true model is 30-fold exponential growth: (a) there is variation in recombination and mutation rates across the genome, (b) there is variation in recombination and mutation rates, and the centromeric region is masked, (c) there is variation in recombination and mutation rates, and short regions resembling repeats are randomly masked across the genome (comprising of 10% of each chromosome), and (d) there is variation in recombination and mutation rates, and the centromere as well as small-sized repeats are randomly masked across the genome. Detailed methods including command can be found here: https://github.com/paruljohri/demographic_inference_with_selection/blob/main/CommandLines/SuppFigure15-20.txt.

**Supp Figure 17:**
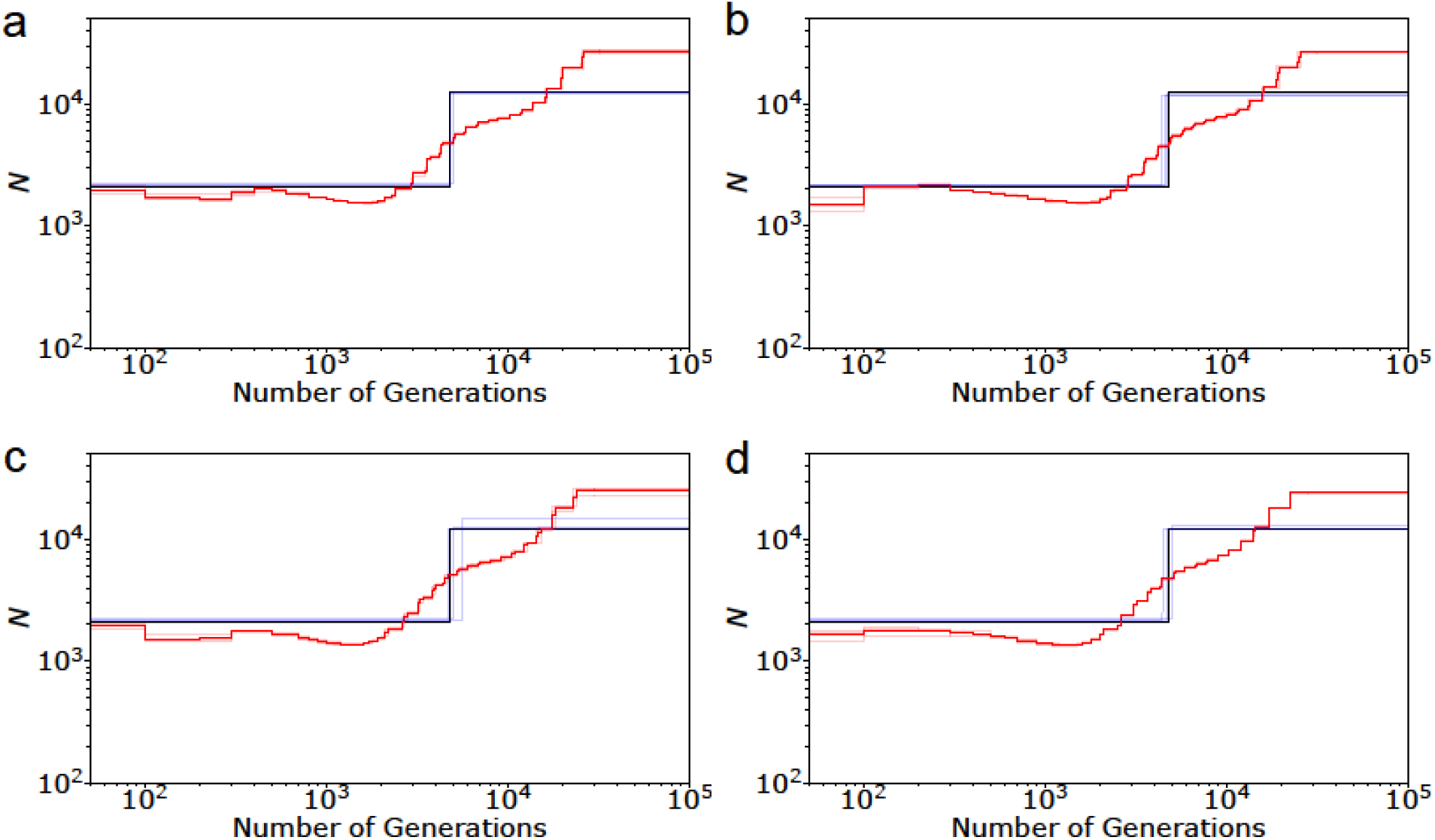
Performance of demographic inference by MSMC (red lines) and *fastsimcoal2* (blue lines) under different scenarios of neutrality, when the true model is 6-fold instantaneous decline: (a) there is variation in recombination and mutation rates across the genome, (b) there is variation in recombination and mutation rates, and the centromeric region is masked, (c) there is variation in recombination sand mutation rates, and short regions resembling repeats are randomly masked across the genome (comprising of 10% of each chromosome), and (d) there is variation in recombination and mutation rates, and the centromere as well as small-sized repeats are randomly masked across the genome. Detailed methods including command lines can be found here: https://github.com/paruljohri/demographic_inference_with_selection/blob/main/CommandLines/SuppFigure15-20.txt.

**Supp Figure 18:**
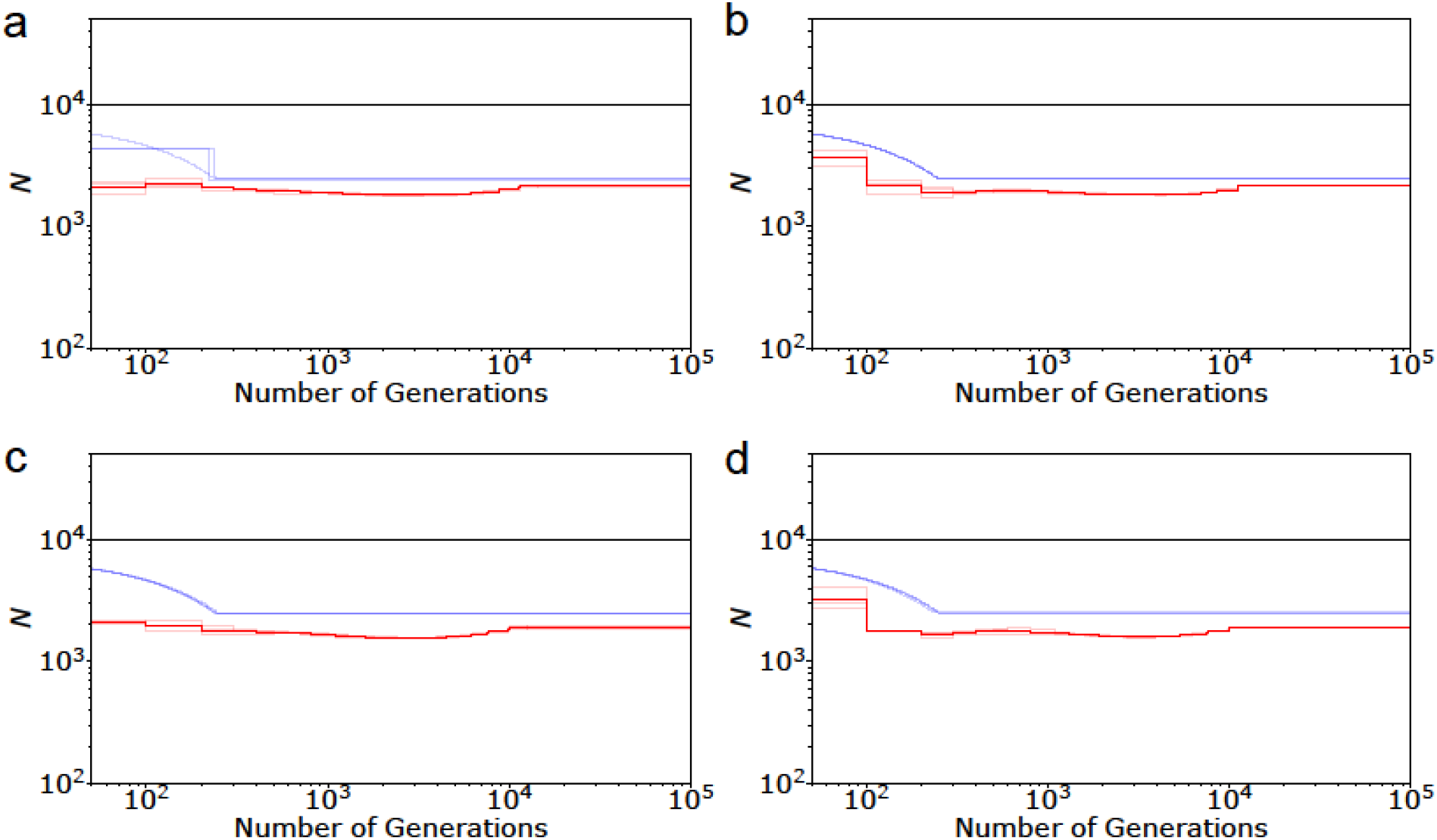
Performance of demographic inference by MSMC (red lines) and *fastsimcoal2* (blue lines) in the presence of background selection, under different scenarios when the true model is equilibrium: (a) there is variation in recombination and mutation rates, (b) there is variation in recombination and mutation rates and the centromeric region is masked, (c) there is variation in recombination and mutation rates, and short regions resembling repeats (comprising 10% of each chromosome) are randomly masked across the genome, and (d) there is variation in recombination and mutation rates, and the centromere as well as small-sized repeats are randomly masked across the genome. Exons comprise of 20% of the genome, experience purifying selection given by DFE4 (*f*_0_ = *f*_1_ = *f*_2_ = *f*_3_ = 0.25), and are masked when performing inference. Detailed methods including command lines can be found here: https://github.com/paruljohri/demographic_inference_with_selection/blob/main/CommandLines/SuppFigure15-20.txt.

**Supp Figure 19:**
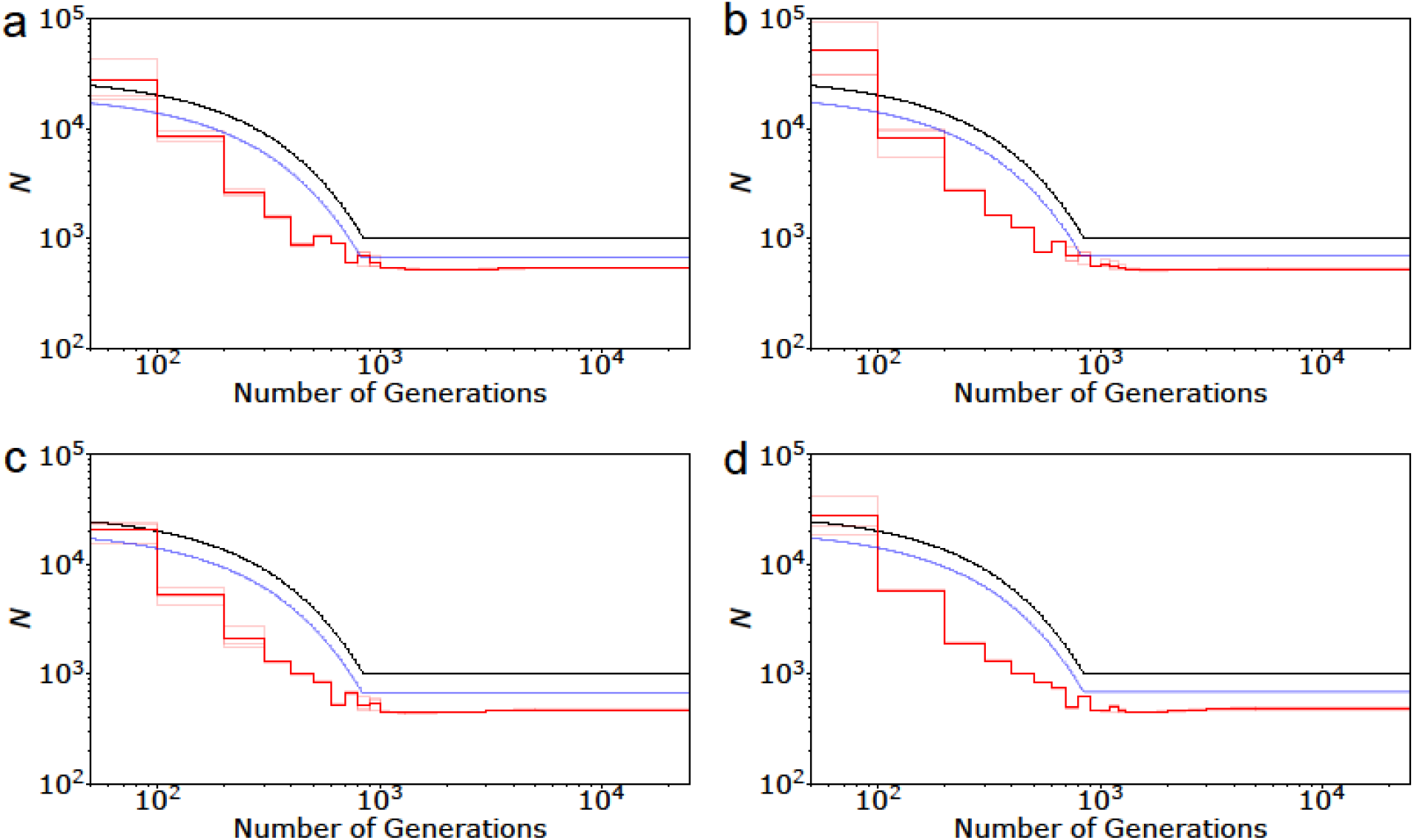
Performance of demographic inference by MSMC (red lines) and *fastsimcoal2* (blue lines) in the presence of background selection, under different scenarios when the true model is 30-fold exponential growth: (a) there is variation in recombination and mutation rates, (b) there is variation in recombination and mutation rates and the centromeric region is masked, (c) there is variation in recombination and mutation rates, and short regions resembling repeats (comprising 10% of each chromosome) are randomly masked across the genome, and (d) there is variation in recombination and mutation rates, and the centromere as well as small-sized repeats are randomly masked across the genome. Exons comprise of 20% of the genome, experience purifying selection given by DFE4 (*f*_0_ = *f*_1_ = *f*_2_ = *f*_3_ = 0.25), and are masked when performing inference. Detailed methods including command lines can be found here: https://github.com/paruljohri/demographic_inference_with_selection/blob/main/CommandLines/SuppFigure15-20.txt.

**Supp Figure 20:**
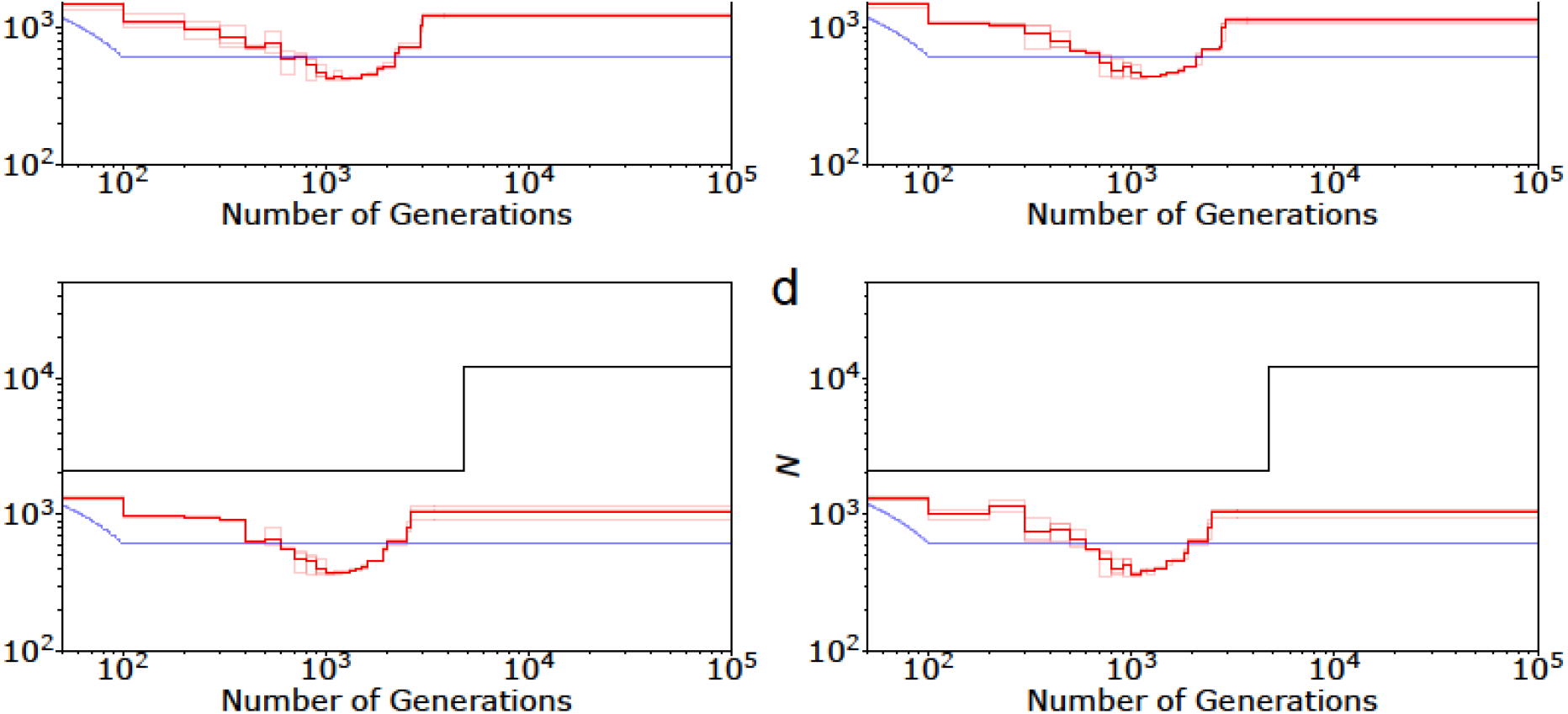
Performance of demographic inference by MSMC (red lines) and *fastsimcoal2* (blue lines) in the presence of background selection, under different scenarios when the true model is a 6-fold instantaneous decline: (a) there is variation in recombination and mutation rates, (b) there is variation in recombination and mutation rates and the centromeric region is masked, (c) there is variation in recombination and mutation rates, and short regions resembling repeats (comprising 10% of each chromosome) are randomly masked across the genome, and (d) there is variation in recombination and mutation rates, and the centromere as well as small-sized repeats are randomly masked across the genome. Exons comprise of 20% of the genome, experience purifying selection given by DFE4 (*f*_0_ = *f*_1_ = *f*_2_ = *f*_3_ = 0.25), and are masked when performing inference. Detailed methods including command lines can be found here: https://github.com/paruljohri/demographic_inference_with_selection/blob/main/CommandLines/SuppFigure15-20.txt.

**Supp Figure 21:**
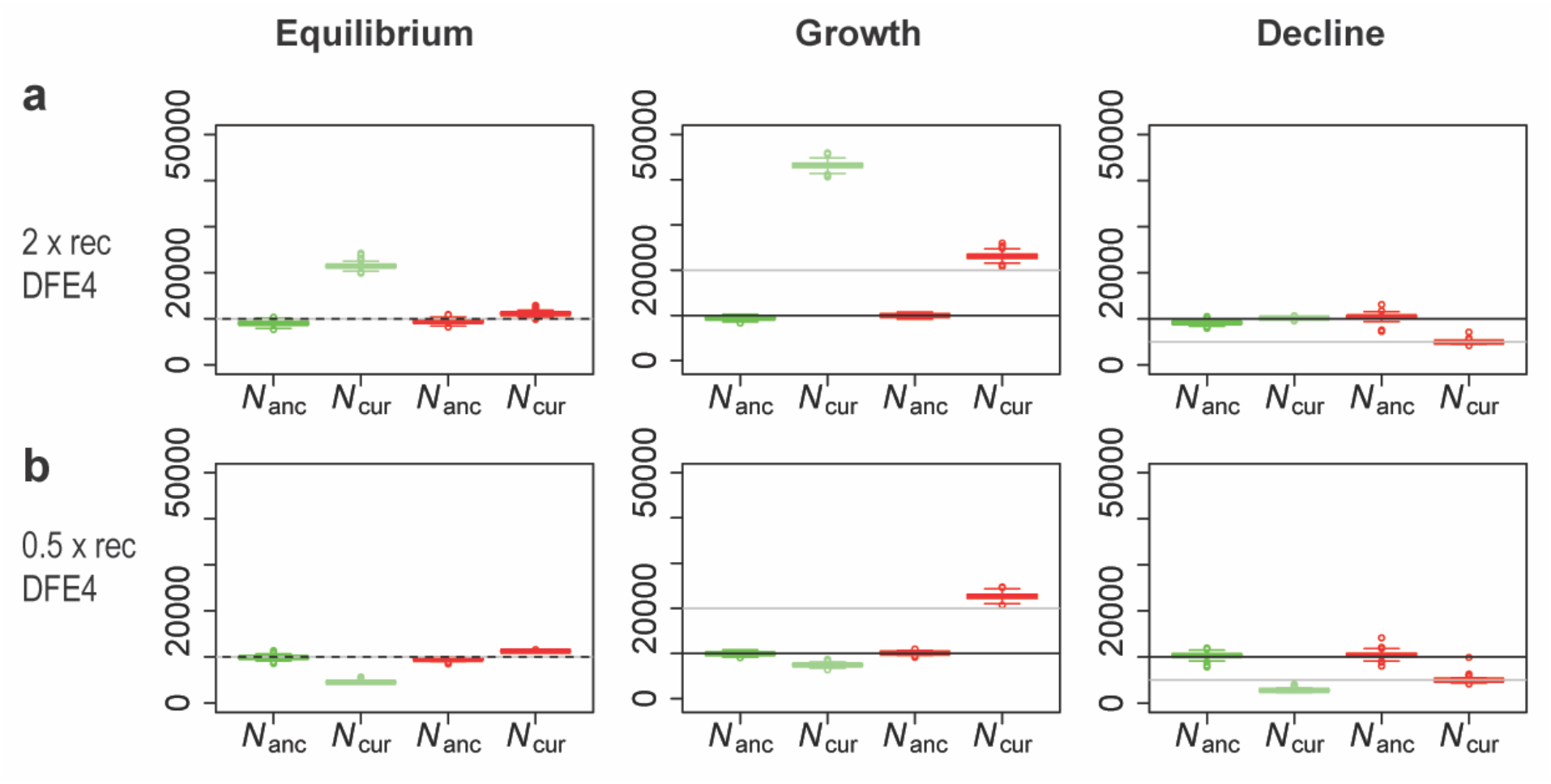
Performance of the ABC method when recombination rate is mis-specified. (a) The true recombination rate is 2-fold higher than that assumed, and (b) the true recombination rate is 2-fold lower than that assumed. Boxplots in green show the posterior estimates when the recombination rate is higher or lower than assumed. For comparison, boxplots in red show the posterior inferred when the corresponding recombination rate is correctly specified. The black line displays the true ancestral population size (*N_anc_*) and the gray line represents the true current population size (*N_cur_*). Detailed methods including command lines can be found here: https://github.com/paruljohri/demographic_inference_with_selection/blob/main/CommandLines/SuppFigure21.txt.

**Supp Figure 22:**
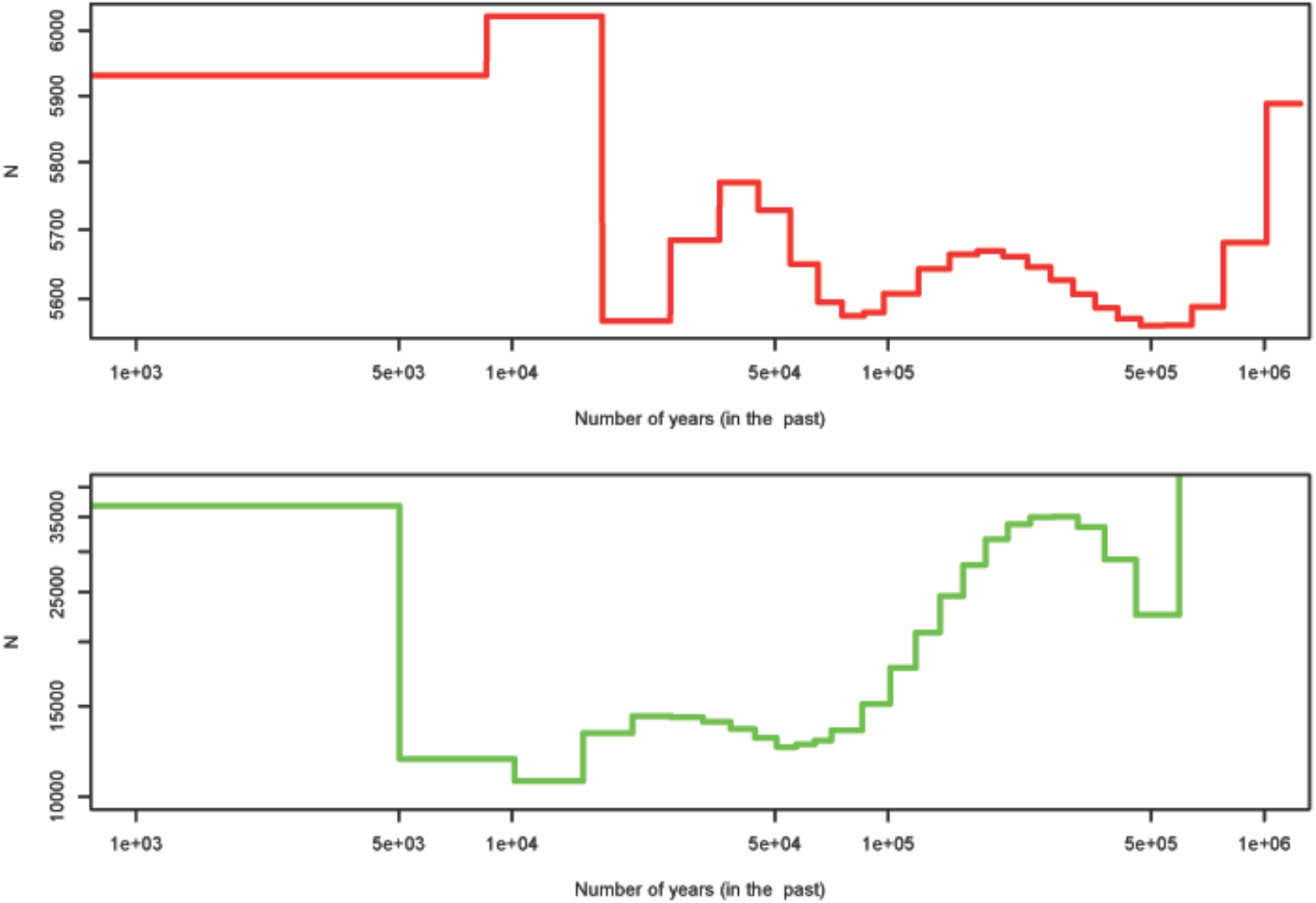
Inference of demographic history by MSMC. Top panel / red line: simulations in which the true model is constant population size, and 50% of new mutations in exons are strongly deleterious with the remainder being neutral, where exons comprise 5% of the genome. Bottom panel / green line: the empirical estimate of population history of the YRI population inferred with MSMC by Schiffels and Durbin (2014). The x-axis is in years (assuming a generation time of 30 years). Note that the y-axes are on different scales, and the magnitude of change observed in the empirical data is considerably larger in the simulated data. Thus, this comparison is only meant to illustrate this common shape taken in MSMC plots (and see similar shapes in, for example, vervets (Warren *et al*. 2015; Figure 4) and passenger pigeons (Hung *et al*. 2014; Figure 2)).

